# Piezo1-induced durotaxis of pancreatic stellate cells depends on TRPC1 and TRPV4 channels

**DOI:** 10.1101/2023.12.22.572956

**Authors:** Ilka Budde, André Schlichting, David Ing, Sandra Schimmelpfennig, Anna Kuntze, Benedikt Fels, Joelle M-J Romac, Sandip M Swain, Rodger A Liddle, Angela Stevens, Albrecht Schwab, Zoltán Pethő

**Affiliations:** Institute of Physiology II, University of Münster, Robert-Koch Str. 27B, 48149, Germany; Institute for Analysis and Numerics, University of Münster, Einsteinstr. 62, 48149, Germany; Gerhard-Domagk-Institute of Pathology, University of Münster; Münster, Germany; Institute of Physiology, University of Lübeck; Lübeck, Germany; Department of Medicine, Duke University, Durham, North Carolina, 27708, USA

**Keywords:** mechanosensation, mechanotransduction, pancreatic cancer, taxis

## Abstract

Pancreatic stellate cells (PSCs) are primarily responsible for producing the stiff tumor tissue in pancreatic ductal adenocarcinoma (PDAC). Thereby, PSCs generate a stiffness gradient between the healthy pancreas and the tumor. This gradient induces durotaxis, a form of directional cell migration driven by differential stiffness. The molecular sensors behind durotaxis are still unclear. To investigate the role of mechanosensitive ion channels in PSC durotaxis, we established a two-dimensional stiffness gradient mimicking PDAC. Using pharmacological and genetic methods, we investigated the role of the ion channels Piezo1, TRPC1, and TRPV4 in PSC durotaxis. We found that PSC migration towards a stiffer substrate is diminished by altering Piezo1 activity. Moreover, disrupting TRPC1 along with TRPV4 abolishes PSC durotaxis even when Piezo1 is functional. Hence, PSC durotaxis is optimal with an intermediary level of mechanosensitive channel activity, which we simulated using a numerically discretized mathematical model. Our findings suggest that mechanosensitive ion channels, particularly Piezo1, detect the mechanical microenvironment to guide PSC migration.

**Summary:** Cells move towards regions with higher stiffness in a process called durotaxis. This study shows that ion channels Piezo1, TRPV4, and TRPC1 are crucial sensors in pancreatic stellate cells. They act together to orchestrate durotaxis of pancreatic stellate cells which may be relevant in pancreatic cancer.

## Introduction

Increased tissue stiffness is characteristic of human solid tumors. Higher stiffness is usually associated with more aggressive tumor growth and, thus, a poorer prognosis (Piersma et al., 2020). Pancreatic ductal adenocarcinoma (PDAC) is a tumor with a particularly stiff – desmoplastic – stroma. A consequence of desmoplasia is an associated increase in tissue stiffness, measurable by atomic force microscopy (AFM) or magnetic resonance elastography. While the healthy pancreas has an average stiffness of 1−2 kPa, PDAC stiffness ranges between 5−10 kPa (Rubiano et al., 2018; Shi et al., 2018). These differences lead to stiffness gradients between the soft, healthy pancreas and the stiff, desmoplastic tumor.

The stromal components in PDAC can account for up to 80% of the tumor mass (Apte et al., 2015). The tumor stroma exerts a malignancy-enhancing effect on PDAC in various ways: (i) pancreatic stellate cells (PSC) produce excess extracellular matrix (ECM) (Pothula et al., 2020); (ii) the associated increase in stiffness further activates and/or maintains the activation of PSCs and (iii) compresses blood vessels; (iv) the reduced blood flow thus abolishes the efficacy of chemotherapy (Apte et al., 2015); (v) carcinoma cells activate PSCs. Reciprocally, PSCs promote cancer cell proliferation, migration, and survival (Apte et al., 2015); and (vi) PSCs may self-activate in an autocrine manner (Pothula et al., 2020; Pethő et al., 2019; Nielsen et al., 2017). Therefore, PSCs and other cancer-associated fibroblasts (CAFs) create and maintain the tumor-promoting microenvironment. Notably, the characteristic mechanical properties of the PDAC microenvironment maintain a positive mechanical feedback loop that further promotes fibrosis (Lachowski et al., 2017).

PSCs sense the stiffness difference between PDAC and healthy pancreatic tissue by migrating toward the stiffer areas (Lachowski et al., 2017). Directional migration along stiffness gradients of the ECM is referred to as durotaxis (Espina et al., 2022). It is also found in other human cell types, including fibroblasts, carcinoma cells, and smooth muscle cells (Lo et al., 2000; DuChez et al., 2019; Evans et al., 2018). While various elements of the migration machinery have been widely documented, the primary mechanosensory transduction process that enables (pancreatic stellate) cells to respond to the stiffness gradients in their environment still needs to be clarified (Lachowski et al., 2018, 2017; DuChez et al., 2019; Janmey et al., 2020; Espina et al., 2022).

Since cells can respond to mechanical stimuli in their environment, they must have an appropriate sensory mechanism. Several potential mechanosensory and mechanotransduction pathways exist. These involve integrins, focal adhesion complexes, G protein-coupled receptors, and mechanosensitive ion channels (Espina et al., 2022). Many mechanosensitive ion channels are Ca^2+^-permeable and are close to focal adhesions (Kobayashi and Sokabe, 2010). They can detect mechanical stress changes and translate them into a Ca^2+^-influx. Thus, Ca^2+^ is a central second messenger that can initiate global cellular responses to a local mechanical event (Nourse and Pathak, 2017). However, the functional relationship between mechanosensation and the initiation of the corresponding cellular response still needs to be understood.

Prominent mechanosensitive ion channels in PSCs include Piezo1 and TRPV4 (Coste et al., 2010; Goswami et al., 2010; Swain et al., 2022). Piezo1 is expressed in PSCs and is involved in cell migration in a pH-dependent manner (Fels et al., 2016; Kuntze et al., 2020). Moreover, Piezo1 and TRPV4 play a role in pressure-induced pancreatitis, which may act as a precursor lesion for PDAC (Swain et al., 2020). In the process of further transducing the mechanical signals, other ion channels expressed in PSCs may also be involved in an indirect manner, such as TRPC1 and K_Ca_3.1 (Storck et al., 2017; Fabian et al., 2012). TRPC1 transduces the pressure-induced activation of PSCs (Fels et al., 2016; Radoslavova et al., 2022). The expression pattern of the ion channels also shows a reciprocal influence: knockout of the TRPC1 channel in PSCs leads to overexpression of the TRPV4 channel (Fels et al., 2016).

Whether mechanosensitive ion channels in PSCs play a role in stiffness gradient perception is currently unexplored. Ion channels are key integration sites for polymodal extracellular stimuli (Pethő et al., 2019). Thus, we hypothesized that they act as a “missing link” between ECM stiffness perception and durotaxis. This work aims to show to what extent mechanosensitive ion channels mediate PSC durotaxis.

## Results

### PSC phenotype is altered in a stiffness-dependent manner

First, we tested whether the phenotype of PSCs differs when they are seeded on soft and stiff matrices reflecting the properties of the healthy pancreas and desmoplastic PDAC, respectively. For this, we created polyacrylamide hydrogels (Table S1) coated with a thin layer of ECM. These hydrogels accurately mimic the substrate stiffness levels of the healthy pancreas (750 Pa) and the increased stiffness seen in PDAC (5 kPa), as well as an even greater stiffness (13.5 kPa) (Fig S1A) (Rubiano et al., 2018; Shi et al., 2018). We seeded murine PSCs onto the hydrogels and observed their two-dimensional phenotype (Fig. 1A) and migratory behavior (Fig. 1B). As observable from the durotaxis plots (Fig. 1B), PSCs have no preferential directionality without a stiffness gradient (Fig. S1B). Moreover, PSCs seeded on a soft 750 Pa substrate migrate substantially more slowly than cells seeded on stiffer substrates (Fig. 1C). PSCs spread more on stiffer substrates, as cells on a 750 Pa hydrogel have ∼10 times less area (Fig. 1D) and fivefold higher circularity (Fig. 1E) compared to PSCs on 5 kPa and 13.5 kPa substrates. Cells seeded onto 5 kPa and 13.5 kPa substrates behave similarly, suggesting that PSC spreading approaches a maximum and cannot be further increased by higher stiffness values.

**Figure 1.**
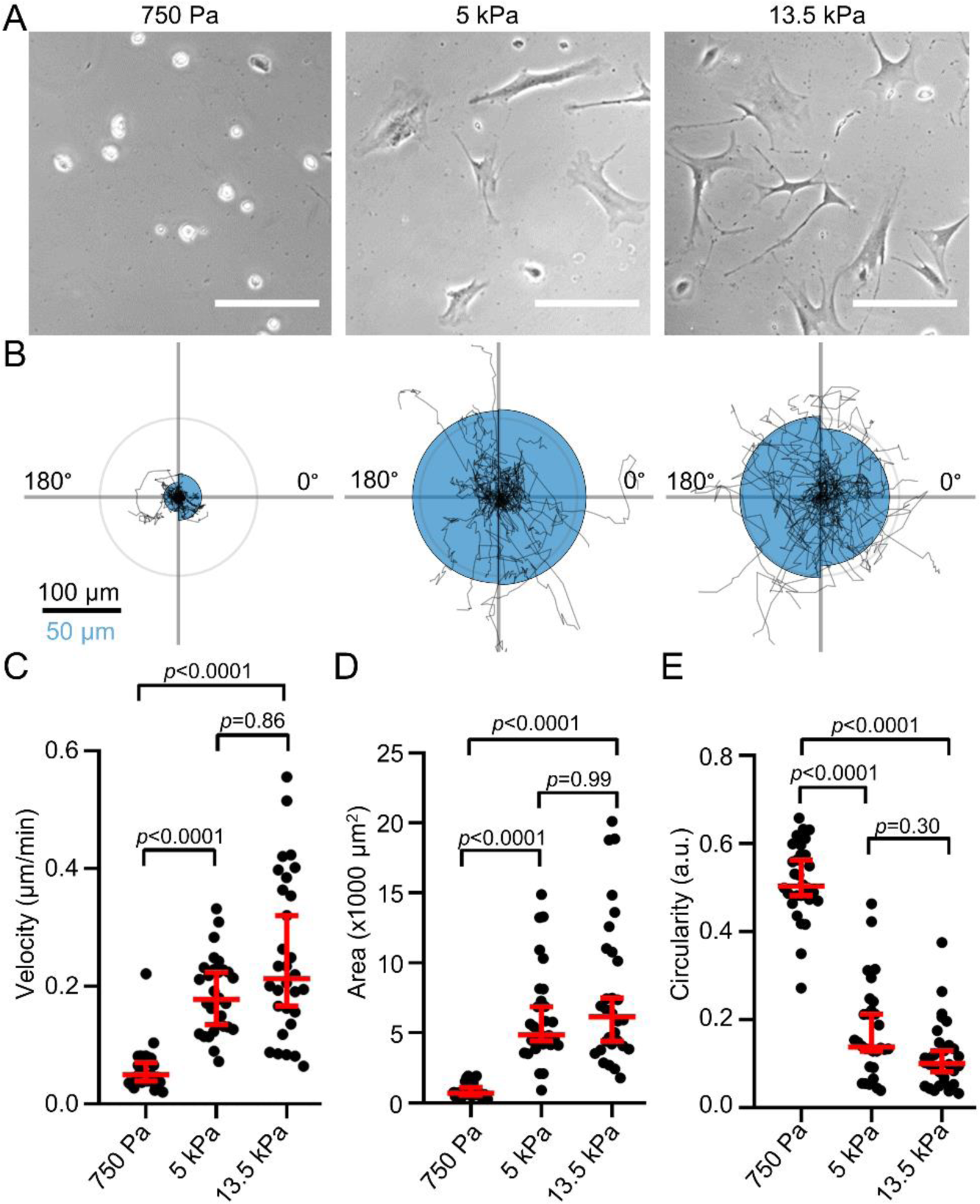
Substrate stiffness affects pancreatic stellate cell morphology and migration. (A) Representative phase-contrast microscopy images of PSCs seeded on substrates with different stiffnesses. Scale bar = 100 µm (B) Durotaxis polar plots depict individual PSC trajectories over 24 h (black lines) on substrates with stiffnesses as indicated in (A). The radius of the blue half circles is proportional to the mean cellular displacement towards 0° and 180°, respectively. Radial lines indicate 0°, 90°, 180°, 270°. The scale bar corresponds to two sizes: 100 µm for the trajectories and 50 µm for the half circles. The radius of the concentric circle is a visual aid for the scale bar. (C-E) Scatter plots depict PSC velocity, area, and circularity on substrates with indicated stiffness (n cells / N experiments = 30/3). Data and statistical comparison in (C), (D), and (E): median±95% CI with Kruskal-Wallis statistical test with Dunn’s post-hoc test.

### PSCs undergo durotaxis under pathophysiologically relevant stiffness conditions

Next, we assessed how PSCs perceive spatial differences in substrate stiffness representing the soft, healthy pancreatic (1−2 kPa) and stiff tumorous tissue (5−10 kPa). Therefore, we established a gradient hydrogel system recapitulating both stiffnesses. We validated the substrate stiffness by AFM (Fig. 2A-C). The AFM measurements show a linear drop in stiffness of ECM-coated gels by ∼5 kPa over a distance of 2 mm in the region of the gradient (Fig. 1B). The stiffest area of the gradient hydrogel is in the upper range of the stiffness values of PDAC, whereas the softest area of the gel is in the upper range of the physiological stiffness of the healthy pancreas (Rubiano et al., 2018; Shi et al., 2018). The gel areas where the photomask was transparent (positions -5 – 1 mm) show a constant stiffness, further underlying the specificity of UV-polymerization with a photomask. To exclude that ECM coating affects hydrogel properties, we systematically investigated the gradient stiffness of the same gels before and after ECM coating (Fig. 2B) as well as before and after the 24 h long cell migration assays (Fig. 3C). Importantly, the stiffness gradient remained constant throughout the whole experimental period.

**Figure 2.**
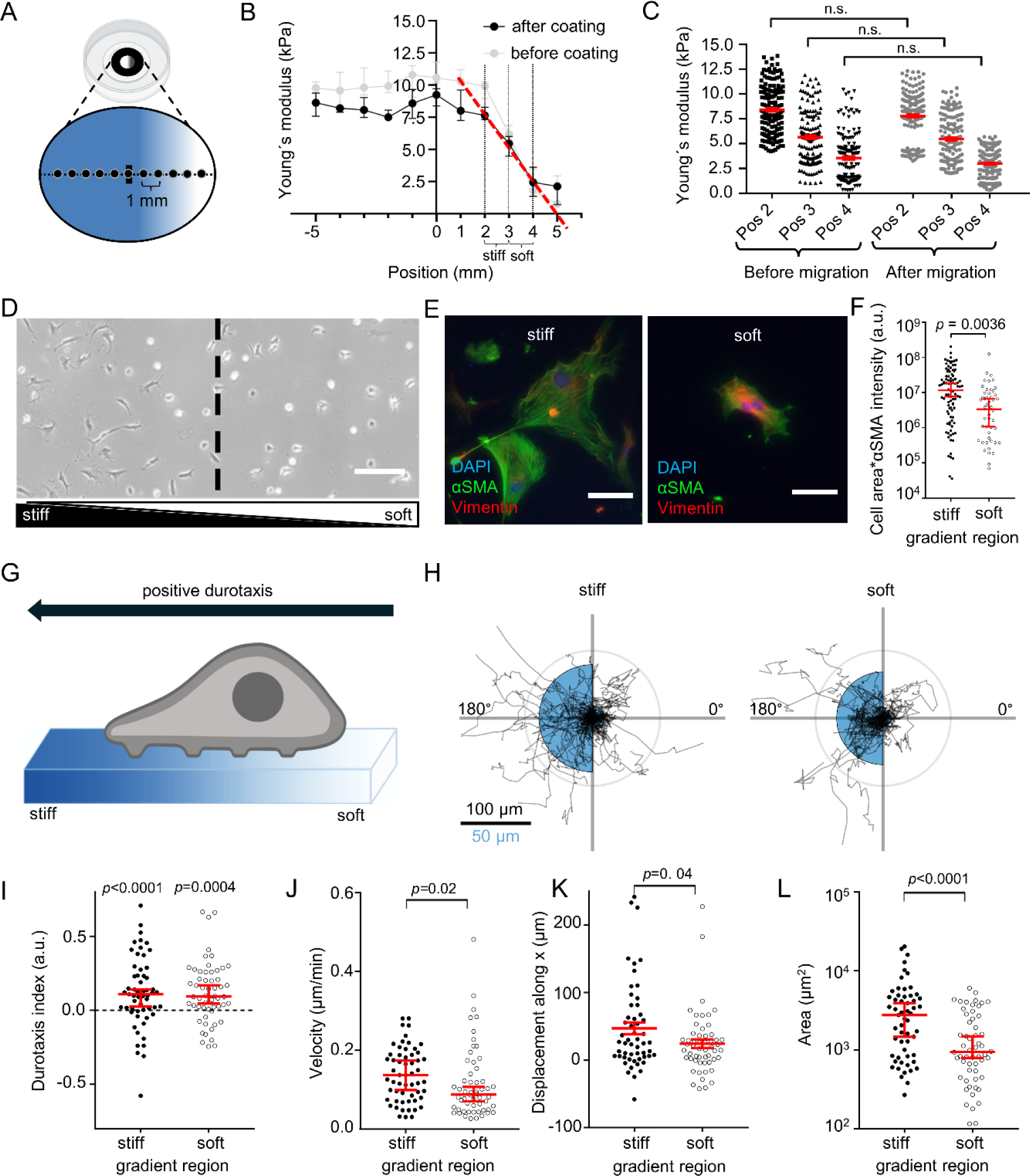
PSCs undergo durotaxis on linear stiffness gradient hydrogels. (A) Experimental setup of durotaxis: glass-bottom dishes are coated with a UV-polymerized hydrogel using a gradient photomask (greyscale form inside the dish). The resulting stiffness gradient of the gel was measured with AFM from stiff (blue) to soft (white), starting from the middle of the gel (horizontal line) in 1 mm steps (black dots on line). (B) Gradient hydrogel stiffness before (grey) and after (black) ECM coating. n points measured / N gels = 40/4. The stiffest area of the gradient hydrogel (position 2 mm, “stiff”) has a median stiffness of 7.6 kPa (95%-CI: 7.3–8.3 kPa), whereas the soft area of the gel (position 4 mm, “soft”), has a median stiffness of 2.4 kPa (1.3–3.6 kPa). Note the linear decay of stiffness in the gradient region after coating (red line, slope = –2.47 kPa/mm). (C) Scatter plots depict gel stiffness measured before (black) and after (gray) cell migration assays. n points measured / N gels = 33/3. (D) Representative phase-contrast image of PSCs on an ECM-coated gradient hydrogel. The dashed line indicates the center of the gradient separating “stiff” and “soft” gradient regions. Scale bar = 250 µm. (E) Representative immunofluorescence images of the myofibroblastic PSC marker αSMA (green), vimentin (red) and DAPI (blue) of PSCs seeded on stiff (left) and soft (right) regions, respectively, from N=4 experiments. Scale bar = 150 µm. (F) Scatter plot of total PSC αSMA fluorescence assessed by multiplying cell area with αSMA fluorescence intensity. n cells measured / N experiments ≥ 51/4. (G) Schematic depiction of “positive” durotaxis. The cells move towards the stiffer side of the gels. (H) Durotaxis polar plots depict individual PSC trajectories over 24 h (black lines). The radius of the blue half circles is proportional to the mean cellular displacement into the directions 0° and 180°, respectively. Radial lines indicate 0°, 90°, 180°, 270°. The scale bar corresponds to two sizes: 100 µm for the trajectories and 50 µm for the half circles. Radius of the concentric circle is a visual aid for the scale bar. (I-L) Scatter plots show the PSC durotaxis index, displacement along the x axis, and velocity, respectively, on stiff (left) and soft (right) areas of the gradient hydrogels. n cells / N experiments = 56/4. Data in (B), (C), (F), and (I-L): median±95% CI. Statistical tests in (C), (F), and (J-L) are Mann-Whitney U-test, and in (I) one-sample Wilcoxon test. Images in (A) and (G) were created with BioRender.com.

**Figure 3.**
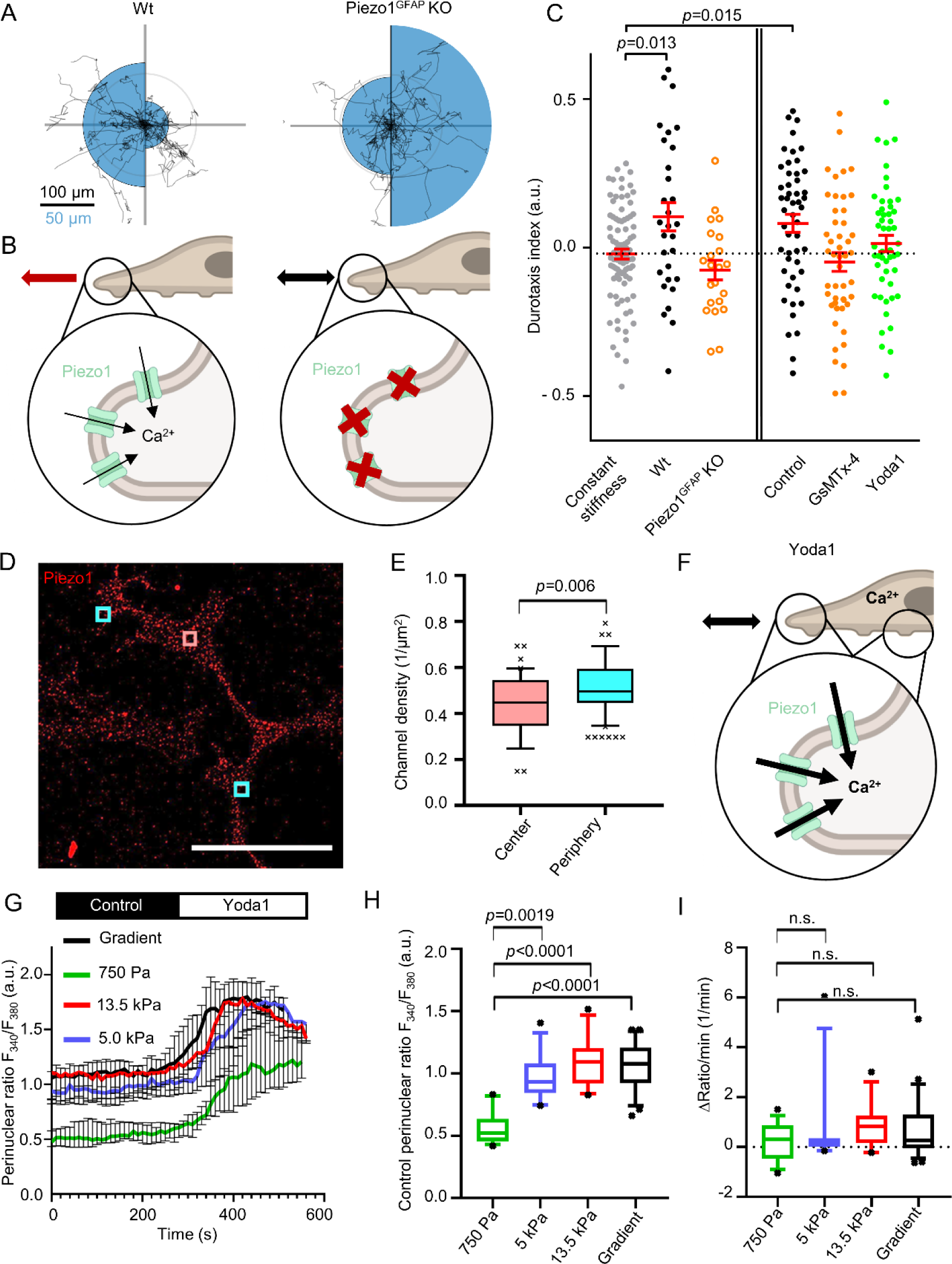
Piezo1 is involved in PSC durotaxis. (A) Durotaxis polar plots of wildtype (Wt, left) and Piezo1^GFAP^ knockout PSCs depict cell trajectories over 24 h (black lines). The radius of the blue half circles is proportional to the mean cellular displacement into the directions 0° and 180°, respectively. Radial lines indicate 0°, 90°, 180°, 270°. The scale bar corresponds to two sizes: 100 µm for the trajectories and 50 µm for the half circles. Radius of the concentric circle is a visual aid for the scale bar. n_Wt_ = 29, n_Piezo1-GFAP_ KO = 22 cells from N=3 experiments. (B) Interpretation of results depicted in (A). In Piezo1^GFAP^ KO PSCs the intracellular Ca^2+^ signal is missing in response to stiffness sensing. Thus, Piezo1^GFAP^ KO PSCs fail to perform durotaxis. (C) Durotaxis indices of Piezo1^GFAP^ KO PSCs (orange) or PSCs treated with 100 nM GsMTx-4 (orange), 5 µM Yoda1 (green), and their respective controls (black) on stiffness gradient hydrogels, compared to PSCs on gels with constant stiffness (grey, dotted line). If cells undergo durotaxis, the *p* value is indicated. n≥22 cells from N≥3 experiments. (D) Representative immunofluorescence image of PSCs stained for Piezo1 (red), from N=3 experiments. Channels were quantified in rectangular regions in the cell center (red) and periphery (cyan). Scale bar = 50 µm. (E) Box and whisker plot shows quantification of Piezo1 channel density in the rectangular regions outlined in (D). n_Center_ =41, n_Periphery_ =82 cells from N=3 experiments (F) Schematic interpretation of how Yoda1 treatment affects durotaxis by clamping Piezo1 activity to its maximal value throughout the cell. Intracellular [Ca^2+^] increases over the whole cell, diminishing the differential stiffness sensing by Piezo1. (G) F_340_/F_380_ ratios were used as a readout for intracellular Ca^2+^ measurements of PSCs. The F_340_/F_380_ ratio is a surrogate of the [Ca^2+^]_i_. After control superfusion (black), PSCs seeded on hydrogels with varying stiffness (750 Pa, 5 kPa, 13.5 kPa, and gradient gels) were superfused with 5µM Yoda1 (white). n ≥ 13 cells from N=3 experiments. (H) Box and whisker plots show quantification of F_340_/F_380_ ratios of PSCs superfused with control solution from experiments detailed in (G). (I) Box and Whisker plots indicate the slope of F_340_/F_380_ ratio upon Yoda1 superfusion, from experiments detailed in (G). Data in (C), (E), and (G-I) are median±95% CI. Boxes in (E), (H) and (I) extend from the 25th to 75th percentiles, whiskers from 10 to 90 percentiles, and points below and above the whiskers are drawn as individual points. Statistical tests are as follows: in (C) one-sample Wilcoxon test; in (E) Mann-Whitney U-test, and in (H-I) Kruskal-Wallis test with Dunn’s post-hoc test. Images in (B) and (E) were created with BioRender.com.

After seeding PSCs onto the hydrogels with a stiffness gradient, we observed that PSC morphology is different in stiff and soft areas of the gradient (Fig. 2D), reminiscent of PSC morphology on stiff and soft hydrogels (Fig. 1A). Next, we assessed the myofibroblastic phenotype of PSCs based on total cellular αSMA (Pethő et al., 2023), derived from αSMA fluorescence intensity multiplied by cell area (Fig. 2D-E). PSCs have more αSMA on the stiffer part of the gradient than on the softer part. Also, cell area and αSMA fluorescence intensity themselves significantly differ between stiff and soft regions (Fig S2A-B).

Next, we monitored the migratory behavior of PSCs over 24h on the ECM-coated gradient hydrogels (Fig. 2G-L). We visualized the functional mechanosensitivity of PSCs using polar plots (Fig. 2H) and assessed the durotaxis index as a readout of the relative migration toward the stiff substrate (Fig. 2I). The PSCs have a positive durotaxis index, both on the stiffer and the softer halves of the gradient. On the stiffer part of the gel, the net displacement along the stiffness gradient (x axis) is higher (Fig. 2K), because the cells generally migrate faster (Fig. 2J). Moreover, PSCs have a larger area (Fig. 2L) on the stiffer regions of the hydrogel. These results are consistent with the observations on the homogeneous gels (Fig. 1C-E): Higher stiffness is associated with increased cell area and velocity.

### PSC durotaxis depends on Piezo1 channel activity

Having demonstrated that PSCs exhibit a stiffness-guided migratory behavior, the following sections focus on better understanding the molecular mechanisms behind the process of durotaxis. Mechanically evoked transient Ca^2+^ spikes elicited by ion channel activity at the anterior cell pole of migrating cells are known to control the directionality of cell movement (Wei et al., 2009; Holt et al., 2021). With this in mind, we performed a proof-of-principle experiment to test whether the mechanosensitive ion channel Piezo1 is involved in durotaxis. We employed a genetic model and used PSCs from wild-type (Piezo1^fl/fl^) mice or from mice that harbor a GFAP-dependent conditional Piezo1 knockout (Fig. 3A). The Piezo1^GFAP^ KO PSCs elicit no Ca^2+^ signal upon Yoda1 stimulation (Fig. 3B, Fig S3A-B)(Romac et al., 2018; Swain et al., 2022). Indeed, Piezo1^GFAP^ KO PSCs do not perform durotaxis on ECM-coated gradient hydrogels, as compared to control PSCs on gels with uniform stiffness. Furthermore, we modulated Piezo1 in PSCs pharmacologically: either inhibiting the channel with 100 nM GsMTx-4 or activating the channel using 5 µM Yoda1 (Fig. 3C, Fig S4). While the control PSCs, treated with 0.1% DMSO, undergo durotaxis, both GsMTx-4- and Yoda1-treated PSCs have a durotaxis index not significantly different from 0 (corresponding to random migration). The data from GsMTx-4- and Piezo1^GFAP^ KO PSCs indicate that the ability of PSCs to dynamically adapt their Piezo1 activity to environmental cues is essential for durotaxis (Fig. 3B).

We further investigated why Piezo1 activation with Yoda1 diminishes durotaxis (Fig. 3C). One possibility is that spatially different substrate stiffness signals to the cell by localized Piezo1 activity. Alternatively, inhomogeneous channel expression could transmit differences in substrate stiffness. To test the spatial heterogeneity of Piezo1 in the plasma membrane of PSCs, we performed Piezo1 immunocytochemistry on the gradient hydrogels. Channel distribution was recorded by counting channels in 5 µm^2^ regions (Fig. 3D). The measured regions were either in the cell periphery or located in the cell center. As Fig. 3E illustrates, the average channel density ranges around ∼0.5 channels / µm^2^, with the cell periphery having a marginally higher channel density than the cell center. This would imply that global Piezo1 activation by Yoda1 elicits similar Ca^2+^ signals throughout the cell that are insufficient for spatial stiffness sensing (Fig. 3F). Measurements of the intracellular Ca^2+^ ([Ca^2+^]_i_) revealed that the initial F_340_/F_380_ ratio is significantly lower in PSCs on a 750 Pa substrate compared to cells on stiffer substrates (Fig. 3H). However, upon superfusing the PSCs with 5 µM Yoda1, the F_340_/F_380_ ratio increased similarly on each substrate, implying a full activation of Piezo1 throughout the plasma membrane (Fig. 3I).

### Piezo1 orchestrates PSC durotaxis together with TRPV4 and TRPC1

There is strong evidence that Piezo1 acts in conjunction with another mechanosensitive ion channel, TRPV4 – which acts as a signal amplifier for pressure sensing – to execute cellular functions (Swain et al., 2020). Thus, we next tested whether TRPV4 is involved in PSC durotaxis. For this purpose, we either applied a pharmacological TRPV4 activator (GSK1016790A; 100 nM) or the TRPV4 inhibitor HC067047 (100 nM). Additionally, we investigated whether the TRPC1 channel mediates durotaxis, as the channel is required for directional migration in response to chemical and mechanical cues (Fabian et al., 2008, 2012). Also, we found this channel to be a transducer of pressure-mediated PSC activation (Radoslavova et al., 2022). For this, we used PSCs isolated from TRPC1-KO mice.

We found that pharmacological activation of TRPV4 impairs durotaxis in PSCs isolated from WT and TRPC1-KO mice (Fig. 4A, Fig S5). This effect is similar to the Piezo1-activation found with 5 µM Yoda1 (Fig. 3C, Fig S3). In contrast, TRPV4 inhibition in PSCs from WT mice, or TRPC1 knockout alone, do not affect durotaxis. However, inhibiting TRPV4 in PSCs from TRPC1-KO mice hampers durotaxis, indicating that Piezo1 channels by themselves are not sufficient to orchestrate durotaxis but rather require TRPV4 and/or TRPC1 channel activity.

**Figure 4.**
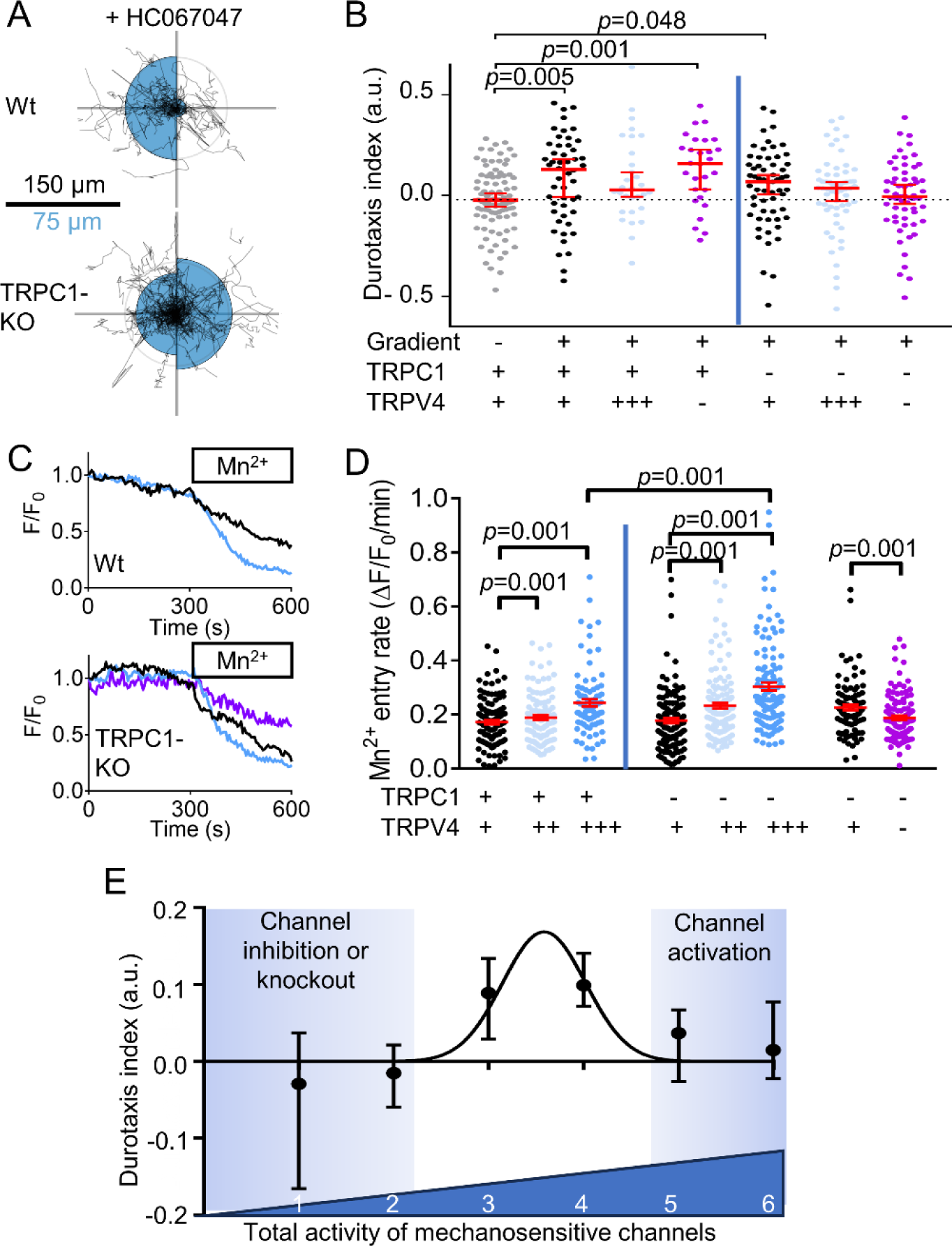
Multiple mechanosensitive channels are necessary for durotaxis. (A) Durotaxis polar plots of wildtype (Wt, top) and TRPC1 knockout (TRPC1-KO, bottom) PSCs depict cell trajectories over 24 h (black lines) under treatment with the TRPV4 inhibitor 100 nM HC067047. The radius of the blue half circles is proportional to the mean cellular displacement into the directions 0° and 180°, respectively. Radial lines indicate 0°, 90°, 180°, 270°. The scale bar corresponds to two sizes: 100 µm for the trajectories and 50 µm for the half circles. Radius of the concentric circle is a visual aid for the scale bar. n_Wt_ = 29, n_Piezo1-GFAP_ KO = 22 cells from N=3 experiments. (B) Durotaxis indices of n wild-type PSCs (TRPC1 +) / N experiments = 28/3 and n PSCs from TRPC1-KO mice (TRPC1 -) / N experiments = 59/3 (left and right from vertical line, respectively) seeded on gradient hydrogels treated with 0.1% DMSO (black), 10 µM TRPV4-activator GSK101489 (TRPV4 +++, light blue), 100 nM HC067047 (TRPV4-, purple). They are compared with PSCs seeded on gels with homogeneous stiffness (grey, dotted line). If cells undergo durotaxis, the *p* value is indicated. (C) Representative graphs showing the relative fluorescence intensity (F/F_0_) of Fura-2 AM loaded PSCs. Under continuous of Mn^2+^ (box), quenching of the Fura-2 signal can be observed under control (black), in the presence of 20 nM GSK101489 (blue) or 2 µM HC067047 (purple). The Mn^2+^ quenching was performed in control cells (grey line) and in pressurized PSCs (black line). (D) Mn^2+^-entry rates of wild-type PSCs and PSCs from TRPC1-KO mice determined under control conditions (TRPV4 +) and in the presence of GSK101489 (20 nM, TRPV4 ++; 100 nM, TRPV4 +++) or 2 µM HC067047 (TRPV4 -). n≥87 cells from N=3 experiments. (E) Durotaxis indices as a function of total mechanosensitive ion channel activity. To estimate total mechanosensitive channel activity, individual channel activity was binned: impaired channel = 0; intermediate activity = 1 for TRPV4 and TRPC1 and 2 for Piezo1; overactivation= 3 for TRPV4 and 4 for Piezo1. A Gaussian curve was fitted over the data. Data in (A) and are median±95% CI. Statistical test in (A) is one-sample Wilcoxon test.

We assumed that the observed effects of TRPV4 and TRPC1 modulation on durotaxis are due to the Ca^2+^ influx mediated by these channels. To mechanistically dissect how channel activity and Ca^2+^ influx are affected by TRPV4 and TRPC1 modulation, we next performed a set of Mn^2+^ quench experiments on PSCs. Here, the Mn^2+^ entry rate is a surrogate of the Ca^2+^ influx into the PSCs. As observable from Fig. 4B, the TRPV4 activator GSK1016790A acts in a dose-dependent manner on PSCs: in WT PSCs 100 nM but not 20 nM GSK1016790A elicit a marked Ca^2+^ influx. Moreover, TRPV4 modulation leads to more pronounced effects in TRPC1-KO PSCs as compared to WT PSCs. In TRPC1-KO PSCs, already 20 nM GSK1016790A are sufficient to increase the Ca^2+^ influx rate, and 100 nM GSK1016790A significantly further amplify Ca^2+^ influx as compared to WT. We also tested whether diminished durotaxis upon of TRPV4 inhibition in TRPC1-KO is due to impaired Ca^2+^ influx. Indeed, we found a decreased Ca^2+^ influx rate in the presence of the TRPV4 inhibitor HC067047 (100 nM) versus the control.

The results above imply that durotaxis depends in a bell-shaped manner on Piezo1 and TRPV4 channel activity. Both fixed loss of the Piezo1 channel function by Piezo1^GFAP^ KO or GsMTx-4 and fixed gain of channel function by Yoda1 result in loss of the ability to migrate towards stiffer areas. We found a similar phenomenon upon TRPV4 activation and inhibition, especially when combined with TRPC1 knockout. We next developed a semiquantitative illustration of the bell-shaped dependence of durotaxis on mechanosensitive ion channel activity. In the control state, the intermediate relative activity of TRPV4 and TRPC1 are illustrated by an arbitrary value of 1. Because of the paramount functional relevance of Piezo1 in PSC durotaxis, its activity was assigned an intermediate value of 2. Inhibition or knock-out of TRPV4, TRPC1 and Piezo1 channels correspond to a value of 0. Pharmacological activation of TRPV4 and Piezo1 channels is represented by the values 3 and 4, respectively. The application of these values allows to score the combined activity of mechanosensitive ion channels to values between 2 and 6: 2 indicates GsMTx-4 treatment, Piezo1 knockout, or dual TRPC1-KO and TRPV4 inhibition, where mechanosensitive channels are substantially impaired; a value of 5 or 6 corresponds to a state when TRPV4 and Piezo1 are pharmacologically clamped in an over-activated state (Fig. 4C, detailed in Fig. S4 and Table S2). Our results emphasize that an intermediate level of mechanosenitive channel activity (value of 3-4) is needed for durotaxis. Whenever the dynamics of mechanosensitive ion channel activity are substantially affected, either by tonic inhibition or overactivation, the cell will not be able to sense (and thus react to) changes in local substrate stiffness. Notably, an intermediate activity of Piezo1 is necessary, but not sufficient for durotaxis. Efficient durotaxis of PSCs requires the additional function of TRPV4 or TRPC1.

### A durotaxis model with ion channels and a bell-shaped mechanosensitivity function

Above, we outlined the hypothesis that an intermediary channel activation is necessary for durotaxis. Here, we support and refine this hypothesis with a mathematical model, a system of partial differential equations. Mathematical models of durotaxis have been published previously (Allena et al., 2016) (Malik and Gerlee, 2019). In (Allena et al., 2016) a cellular Potts model is analyzed for a single cell moving over a flat substrate with variable stiffness. The cell is described by several lattice sites. Cell dynamics result from an iterative stochastic reduction of the energy of the whole system, and a modified Metropolis method for Monte Carlo-Boltzmann dynamics is employed. In (Malik and Gerlee, 2019) a stochastic model is introduced where the cell moves by updating its adhesion sites at random times. The rate of updating is determined by the local stiffness of the substrate. From this, a model of partial differential equations of advection-diffusion type is formally derived, where the advective velocity relates to the stiffness, like in the mathematical model we suggest below. In our case, however, we additionally couple to channel dynamics. The advection-diffusion equation in (Malik and Gerlee, 2019) is numerically solved by a Crank-Nicolson finite difference scheme. How to formally obtain advection terms for taxis terms like in (Malik and Gerlee, 2019) was also discussed in (Othmer and Stevens, 2006).

For our mathematical model *u = u(x,t)* denotes the probability density of the cell in space and time, and *ρ* = *ρ*(*x*, *t*), *η* = *η*(*x*, *t*) the densities of the open and closed channels of the cell, respectively. The time-space dynamics of the cell depend on the relative density 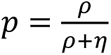 of open channels and is given by

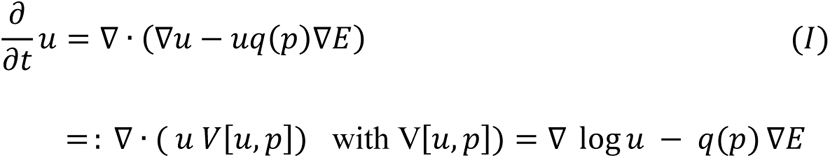

In (I) the first term on the right-hand side accounts for random motion of the cell, and the second term for its tendency to move into the direction of higher stiffness of the substratum. The second line of the formula is a reformulation convenient for the numerics (see details in Numerics). Here *E* denotes the elastic modulus of the substratum, i.e. it is larger for a stiffer material. The function *q* accounts for the strength of the cell to sense the gradient of *E* due to the relative number of open versus closed channels *p*. And it is assumed to be bell-shaped – as suggested by data in Fig. 4 – with a maximum at *p* = 0.5, i.e. *ρ* = *η*, and minimal values at *p* = 0, *p* = 1, i.e. *ρ* = 0, *η* = 1 and *ρ* = 1, *η* = 0. See Fig. 5A-B for the one-dimensional situation on the space interval [0,1]. The dynamics of open and closed channels are given by

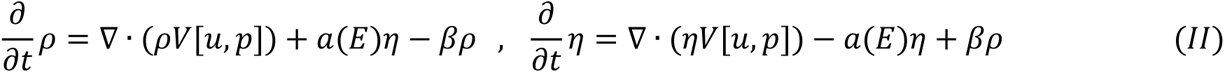

**Figure 5.**
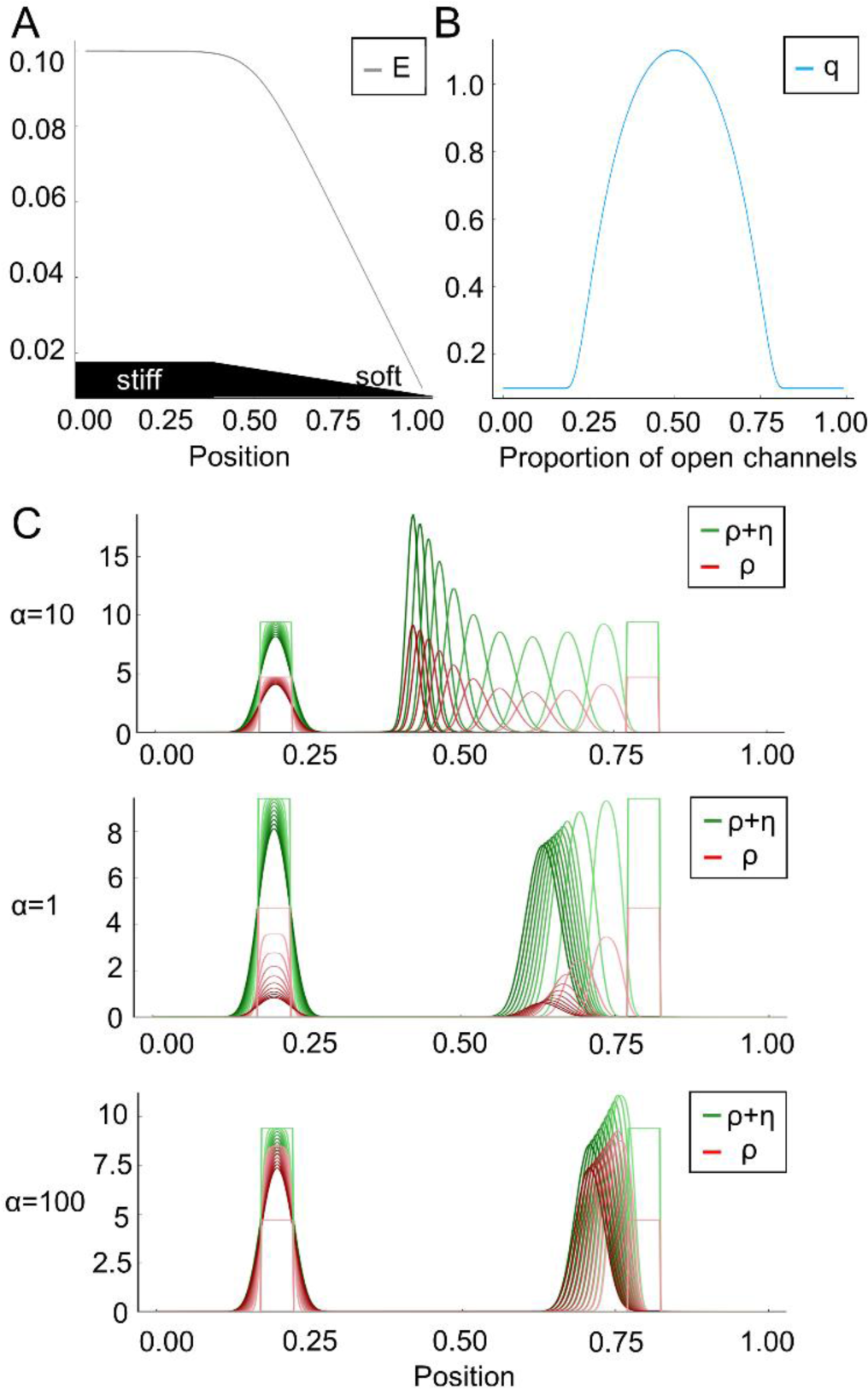
The mathematical model supports the hypothesis of the dependence of durotaxis on ion channel function. (A) *E* denotes the elastic modulus of the substratum, i.e. it is larger for a stiffer material. (B) The bell-shaped function *q* accounts for the strength of the cell to sense the gradient of *E* due to the relative number of open versus closed channels. (C) The diagram depicts the velocity of the probability density of the cell (green) in dependence of the channel opening rate *α*. Red is the density of open channels. *α* = 10 corresponds to an intermediate channel activity, whereas *α* = 1 and *α* = 100 indicate an impaired or overactivated channel function, respectively. The closing rate is *β*=1. For a comparision initially two cells (green rectangles) are located at around 0.2 (stiffer region) and 0.8 (less stiff region). The cell located on the right moves from right to left, i.e. towards the stiffer region. The cell on the left is already located in the stiffer region and changes only slowly.The time frames start from light green/red towards darker green/red. These snapshots are equidistant in time. PSCs do not move towards the stiffer region when the mechanosensitive channels are closed (α = 1) or overactivated (α = 100).

i.e. the channels are located on the cell and thus move with the cell. Further *a* = *a*(*E*) is the opening rate of the channels, which depends on the elastic modulus *E*, and the constant *β* is the closing rate. All partial differential equations are analyzed on a bounded domain with no-flux boundary conditions. Mass of *u* and of *ρ* + *η* is conserved. Further, if initially, i.e. at time *t* = 0, *u*(0, *x*) = *C*(*ρ*(0, *x*) + *η*(0, *x*)) for some constant C > 0, then this feature will be preserved, i.e. *u*(*t*, *x*) = *C*(*ρ*(*t*, *x*) + *η*(*t*, *x*)) for all t > 0. This means that any chosen total initial distribution of channels in the model is preserved over time. In particular we have *V*[*u*, *p*] = *V*[*ρ* + *η*, *p*] in (I) due to the logarithm and the gradient. Once the constant C is fixed, it is sufficient to solve system (II) which can be reformulated in the self-consistent form

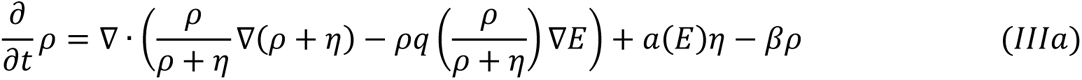

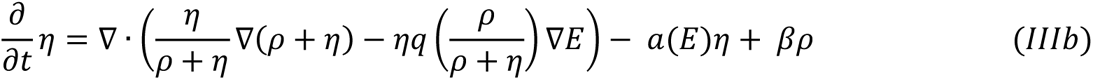

Now we set the opening rate of the channels proportional to the elastic module, i.e. *a*(*E*) = *αE* for *α* > 0. The closing rate *β* is independent of *E*. We consider a one-dimensional spatial setting in the numerical simulations of this mathematical model. Without loss of generality, the strength of random motion is fixed to one in formula (I). When *α* = 10 the simulation results in intermediate mechanosensitive channel activity (Fig. 5C) and recapitulates that cells persist towards the stiffer substrate. Intriguingly, while moving towards the stiffer substrate, the density of open channels increases in time. In contrast, the condition *α* = 1 recapitulates GsMTx-4-treatment, Piezo1^GFAP^ KO PSCs, or TRPC1 KO together with TRPV4 inhibition, where mechanosensitive ion channel function is diminished. Here, impaired channel activity further hampers durotaxis on stiffer substrates over the course of the simulation. A similar inhibition of durotaxis is apparent in the condition *α* = 100 which corresponds to channel overactivation as in the presence of Yoda-1. Together, these simulations show that PSC durotaxis can be effectively modeled via a bell-shaped relationship between mechanosensitive ion channel activity and durotaxis.

## Discussion

In this study, we provide a mechanistic link between ion channels and durotaxis of pancreatic stellate cells. PSC durotaxis is diminished by perturbing ion channels involved in mechanosensing and mechanotransduction (Figs. 3-4). Our main result is that Piezo1 is essential for the durotaxis of PSCs (Fig. 3), but that it depends on the cooperation with TRPC1 and TRPV4 channels (Fig. 4). Overstimulation as well as inhibition of ion channels reduce durotaxis. Our mathematical model supports the conclusion that durotaxis of PSCs is optimal with an intermediate level of dynamically fluctuating activity of mechanosensitive channels (Fig. 5). Stated differently, durotaxis depends on mechanosensitive ion channel activity in a bell-shaped manner. Similarly to channel inhibition, pharmacological activation of Piezo1 using Yoda1 (Syeda et al., 2015) as well as that of TRPV4 by GSK1016790A (Thorneloe et al., 2008) lead to a marked Ca^2+^ influx and inhibit durotaxis (Figs. 3-4). Such bell-shaped relations between migratory behavior and ion transport are also known for other transport proteins such as the Na^+^/H^+^ exchanger (Stock et al., 2005) and K_Ca_3.1 channels (Schwab et al., 1999). Our mathematical model also indicates that PSCs will accumulate on the stiffer substrate over time. These results suggest that PSCs will not migrate out of the stiff tumor tissue. Hence, durotaxis mediated by the mechanosensitive channels constitutes a feed forward mechanism contributing to PSC accumulation and consequent PDAC fibrosis. This adds to the mechanical activation that is also modulated by ion channels involved in mechanosignaling such as TRPC1 (Radoslavova et al., 2022). One strength of our study is that we used substrate rigidities and gradients with pathophysiological relevance. By using a two-dimensional stiffness gradient in our biological system, we were able to monitor every gradient hydrogel with AFM. We found that UV-polymerization using a photomask produced a reproducible, linear stiffness gradient of 2.5 kPa/mm (Fig. 2B). The length of a PSC is ∼50-100 µm, depending on the underlying substrate. This means that a PSC can sense a gradient of <250 Pa between the cell rear and front, on a substrate having a 2–8 kPa stiffness, which underlines the sensitivity of the mechanosensitive cellular network. The durotaxis index is similar on the soft and the stiff part of the gradient gel (Fig. 2I), i.e. the directed and undirected proportion of the cell migration are more or less the same, although cells migrate with a different velocity along the stiffness gradient (Fig. 2J & 2K). This is reflected in the one-dimensional mathematical model by the specific choice of the channel opening rate being proportional to *E*. It likely indicates that the stiffness difference between cell rear and front is more important for durotaxis than the absolute stiffness value. Further studies will outline how these results are applicable to a confined, three-dimensional PDAC tissue.

Our study focuses on substrate stiffness and ignores the viscous properties of the underlying substrate. Notably, it has been shown that elevated viscosity induces a TRPV4-mediated mechanical memory through transcriptional regulation via the YAP-TAZ pathway (Bera et al., 2022). Indeed, YAP1 undergoes nuclear translocation in PSCs on a stiff compared to a soft substrate, resulting in a more myofibroblastic phenotype and increased αSMA expression (Pethő et al., 2023). Hence, it is likely that the tissue viscosity plays an additional role in PSC migration and durotaxis in our experimental system and even more so in the complex PDAC tissue.

Our data suggest that durotaxis relies on the localized activity of mechanosensitive ion channels (Fig. 3). The intracellular calcium concentration of PSCs is influenced by the stiffness of their surrounding ECM. In light of the fact that Piezo1 and TRPV4 have enhanced activity in the vicinity of focal adhesions (Ellefsen et al., 2019; Matthews et al., 2010), a local channel activation is expected in the cell regions adjacent to the stiffer matrix regions of the gradient gel. Therefore, an uncoordinated activation of mechanosensitive ion channels, as provoked using the Piezo1-activator Yoda-1 (Fig. 3G-I) or the TRPV4-activator GSK1016790A (Fig. 4B), would likely prevent durotaxis by disrupting spatiotemporally fine-tuned Ca^2+^ signaling and hence cell polarization. In our system, we could not experimentally test this due to technical limitations, namely due to the thickness and optical properties of the coated polyacrylamide gel. However, it can be assumed that PSCs, like other fibroblasts, actively detect the strength of the surrounding ECM by traction (Sun, 2021). If they are subjected to a stiffness gradient, traction on the stiffer regions will induce less deformation of the substrate. This lower compliance results in a higher membrane tension of the lipid membrane of the cell, which is detected by the Piezo1 channel. According to the force-through-lipid model this provides a suitable opening stimulus for the ion channel (Kobayashi and Sokabe, 2010; Syeda et al., 2016). There is local Piezo1 and TRPV4 activation and a regional influx of Ca^2+^ into the cell, resulting in a stronger pulling than on the soft part of the cell.

Piezo1 and TRPV4 channels act together in pressure-induced pancreatitis (Swain et al., 2022). Similarly, during durotaxis, primary mechanosensing could occur through Piezo1 but be translated into a sufficient cellular response only by subsequent activation of TRPV4. TRPC1 function in durotaxis mechanotransduction supports our previous findings, where we found that TRPC1 perpetuates pressure-mediated PSC activation, as it is required for increased αSMA expression (Radoslavova et al., 2022). Thus, we assume that TRPC1 acts as a transducer of mechanical stimuli, as the intrinsic mechanosensitivity of TRPC1 has not been clearly demonstrated (Nikolaev et al., 2019; Maroto et al., 2005; Gottlieb et al., 2008). Moreover, functional expression of TRPV4 is linked to the presence of TRPC1: TRPV4 is functionally upregulated in TRPC1 KO PSCs, as demonstrated by our Mn^2+^-quench experiments (Fig. 4B). This is well in line with our previous findings pointing out an increased TRPV4 mRNA expression in TRPC1-KO PSCs (Fels et al., 2016).

We argue that the functional interplay between Piezo1, TRPV4, and TRPC1 may have a pathophysiological relevance in PDAC and other tumor entities. A major driver of PDAC progression is the dense, collagen-rich stroma (Pothula et al., 2020). For other tumors, a correlation between elevated tissue stiffness, increased mechanosignaling, and tumor malignancy could be demonstrated (Acerbi et al., 2015). Analogously, for PDAC, a positive mechanical feedback loop involving stiffness-guided activation and migration of PSCs has been postulated (Lachowski et al., 2017; Pethő et al., 2019). Indeed, PSCs spread more on substrates with higher stiffness, leading to a more effective cell migration with a higher velocity (Fig. 1). When introducing a stiffness gradient, PSCs migrate towards a higher stiffness, where they acquire a higher αSMA quantity (Fig. 2). The morphological transition and increase in αSMA point toward an increase in the myofibroblastic machinery, which can explain the higher migration velocity on the stiffer part of the gradient hydrogel (Fig. 2G). In summary, PSC durotaxis relies on the interaction between multiple mechanosensitive ion channels. These results support the idea that the consequences of durotaxis should be further explored in diseases such as PDAC to better understand tumor fibrosis and progression.

## Materials and Methods

### Laboratory animals

The isolation of stellate cells from mouse pancreas has been reported to the Landesamt für Natur, Umwelt und Verbraucherschutz Nordrhein-Westfalen in Recklinghausen under the following number: 84-02.05.50.15.010. If not specified, 8- to 15-week-old wild-type mice (C57 BL/6J) were used for the experiments. In addition, we used pancreatic stellate cells from mice with a global TRPC1 KO and conditional stellate cell-specific Piezo1 knockout (Piezo1^GFAP^ KO) (Dietrich et al., 2007; Swain et al., 2022). PSC isolation from Piezo1^GFAP^ KO mice was performed with approval from the IACUC and IRB of Duke University (protocol number Pro00035974).

### Cells and culture conditions

PSCs were isolated from the pancreas as described before (Kuntze et al., 2020). Briefly, after dissection of the pancreas, the organ was washed in GBSS, then cut into pieces and digested using collagenase P (1 mg/ml) in GBSS for 20 min at 37°C in a shaker. After washing and centrifugation at 300 x g at RT for 8 min, the pelleted pancreas was reconstituted in DMEM/Ham’s-F12 medium. The suspension was seeded dropwise onto FCS-coated cell culture dishes. After 2 h incubation at 37°C and 5% CO_2_, non-adherent cells were washed away by multiple washing steps. Cell removal was validated under the microscope and using fluorescence microscopy. PSCs from Piezo1^GFAP^ KO mice were isolated in a comparable manner, but were plated on a thin-layered Matrigel-coated glass bottom culture plate (MatTek, P35G-0-14-C) (Swain et al., 2022). PSCs in passage 1 were used for all subsequent experiments.

### Polyacryamide hydrogels

Uniform polyacrylamide gels were created using chemical polymerization (Rheinlaender et al., 2020). The polyacrylamide solution contained acrylamide (40 %) (AppliChem GmbH, Germany), bisacrylamide (2 %) (Carl Roth, Germany), hydroxyacrylamide (100 %) (Sigma-Aldrich Chemie, Germany) and PBS. Gel stiffness was tuned by varying the concentrations of the polymer components (acrylamide, bisacrylamide and hydroxyacrylamide) (Supporting Table S1). The solution was ultrasonically degassed for 30 minutes and hereafter, the polymerization was started by adding 0.003 % N,N,N′,N′-tetramethylethylenediamine (TEMED) and 1 % ammonium persulfate (APS) (Sigma-Aldrich Chemie, Germany). 8 µl of the final polyacrylamide mix were pipetted on 35 mm glass bottom dishes (Cell E&G LLC VWR, USA) which were pretreated with 0.1 M NaOH, 200 µl APTMS and 0.5 % glutaraldehyde (SERVA Electrophoresis, Germany). A RainX (ITW Global Brands, USA) coated coverslip was lowered onto the solution and the glass bottom dish was filled with PBS. The coverslips were left overnight and then got removed. The gels were then washed twice with sterile PBS and stored at 4 °C.

For creating hydrogels containing a stiffness gradient, we irradiated a polyacrylamide solution containing the photoinitiator 2-hydroxy-4′-(2-hydroxyethoxy)-2-methylpropiophenone (Irgacure 2959) with UV-light. We controlled the amount of UV-light reaching the gel during the process of polymerization by placing a photomask with a linear opacity gradient between the light source and the gel mix. Thereby, the completeness of the gel formation and hence the stiffness were tuned. The protocol is based on the work of Sheth et al. and Tse and Engler (Sheth et al., 2017; Tse and Engler, 2010). The solution was ultrasonically degassed for 30 minutes and afterwards, the photoinitiator Irgacure 2959 was added. 25 µl of the mix were transferred onto glass bottom dishes pretreated with 0.1 M NaOH and silanized with 200 µl APTMS. A RainX coated coverslip was lowered onto the solution and dishes were then placed on the sample tray of the ChemiDoc™ Touch Imaging System (Bio-Rad, USA). The mask was placed between the dishes and the UV-light source. After 720 seconds of illumination time the gels were immediately treated with PBS and the coverslips were removed.

### Atomic force microscopy

Polyacrylamide hydrogel stiffness and stiffness gradients were quantified using atomic force microscopy (AFM). The Nanowizard 3 NanoOptics AFM system from Bruker Nano GmbH (Berlin, Germany) and the associated SPM software were used for this purpose. Only cantilevers from Novascan Technologies, Inc. (Boone, United States) with spherical tips and a tip diameter of 10 µm were used. According to the manufacturer, the spring constants were between 0.03 and 0.04 N/m and were determined at the beginning of the measurement using the thermal noise method (Cook et al., 2006). The sensitivity calibration of the cantilever was also performed at the beginning of each measurement. For this purpose, a test measurement was made on glass. The sensitivity of the cantilever was calculated from the slope of the linear section of the curve of the deflection signal. All measurements were performed at room temperature and in 2.5 ml sterile PBS. Gel stiffness was determined in 1 mm intervals along the x and y axes. This ensured that the desired stiffness gradient was present in the 2×2 mm area selected for the subsequent tests. Five force-distance curves were generated for each measurement position. 24 h after hydrogel coating (see below), the stiffness was measured in ten further points to verify gradient integrity.

### Hydrogel coating and cell seeding

All hydrogels were coated with a collagen matrix resembling the ECM of the pancreas, containing laminin, fibronectin, and collagens I, III, IV, as previously described (Nielsen et al., 2017; Kuntze et al., 2020). ECM proteins were covalently linked to the gel using Sulfosuccinimidyl-6-(4’-azido-2’-nitrophenylamino)hexanoate (sulfo-SANPAH), a photoreactive heterobifunctional crosslinker. After 15 minutes of irradiation with a wavelength of 302 nm, the gels were washed twice with sterile PBS. After repeating the irradiation and the washing process, 30 µl of the ECM premix were pipetted onto the gel. After a two-hour incubation period at 37 °C, unbound matrix molecules were removed by washing with PBS.

### Measurement of cell migration

Cell migration time-lapse images were obtained using the Axiovert 40C phase-contrast microscope from Carl Zeiss AG (Oberkochen, Germany). The acquisition was positioned on the 4 mm^2^ areas previously measured by AFM. Cell migration was recorded for 24 hours at a time interval of 30 minutes.

Shortly before the start of the experiment, the RPMI medium was replaced by a HEPES-buffered medium. If necessary, the medium of the wild-type PSCs was supplemented with the Piezo1 activator Yoda1 (5 µM), the inhibitor of cation-selective mechanosensitive ion channels GsMTx-4 (100 nM), the TRPV4 agonist GSK1016790A (100 nM) or the TRPV4 inhibitor HC067047 (100 nM) (Syeda et al., 2015; Bae et al., 2011; Thorneloe et al., 2008; Everaerts et al., 2010). TRPC1-KO cells were also treated with the latter two channel modulators (GSK1016790A and HC067047). All listed substances were dissolved in DMSO, therefore 0.1% DMSO was added to the medium of the associated control experiments. Migration videos were analyzed using Amira-Avizo Software 2019.2 (Thermo Fisher Scientific, Waltham, Massachusetts, United States) and by using the Matlab program (MathWorks, Inc., Natick, Massachusetts, United States). Cell outlines were manually segmented in Amira. Subsequently, Matlab calculated the cell perimeter, cell centroid, and cell area. From these data, we calculated further parameters. Cell circularity is quantified by equation

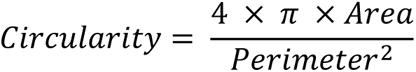

and cell movement parameters were velocity (µm/min; cell centroid displacement as function of time); and durotaxis index, which is defined by:

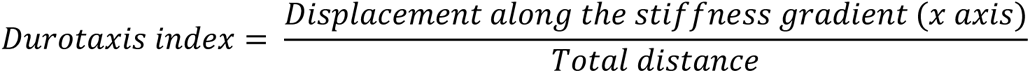

For depicting durotaxis, we developed durotaxis plots using MatLab. For these plots, cell migration data was converted to polar coordinates and individual trajectories were visualized. To explore directional bias, the data was segmented into π radian increments, and the average magnitude of cell movement in each segment was calculated. These averages are then plotted as a qualitative polar histogram. The trajectories and histogram were then overlaid, offering a combined view of individual migration patterns and general directional tendencies.

### Immunofluorescence staining

For α-SMA and vimentin immunostaining of pancreatic stellate cells, gel-containing glass-bottom dishes were fixed with ice-cold methanol after the migration experiments. α-SMA was used as a marker for cell activation, i.e., myofibroblastic phenotype, whereas vimentin was used as a marker to verify the quality of PSC isolation: vimentin positive PSCs were always the overwhelming majority of the cell population, compared to vimentin negative cells of epithelial origin and immune cells. After washing twice with PBS, samples were blocked using 10% FCS in PBS at 4°C for 1 h. Subsequently, primary antibodies against αSMA (A2547, RRID: AB_476701, 1:200, Sigma-Aldrich, Merck KGaA), vimentin (10366-1-AP, RRID: AB_2273020, 1:500, Proteintech) were added to the gels at 4°C for 2 h. After washing three times with PBS, Alexa Fluor 488–conjugated secondary antibodies against mouse (Invitrogen, 1:500) and Cy3-conjugated antibodies against rabbit (Invitrogen, 1:500) and DAPI were added at 4°C for 20 min. An Axio Observer Z.1 inverted microscope from Carl Zeiss AG (Oberkochen, Germany) equipped with a CMOS camera were used to acquire the immunofluorescence images.

For Piezo1 channel staining (Waschk et al., 2011), the gels were washed with PBS then blocked with 1% BSA at 4°C for 1h. Primary antibody against Piezo1 (15939-1-AP, RRID: AB_2231460, 1:200, Proteintech) was added at 4°C for 2 h. Subsequently, the cells were fixed using 3.5% PFA at 4°C for 20 min. Next, secondary antibody and DAPI were added at 4°C for 20 min. Piezo1 fluorescence was recorded using a DMI 6000 Confocal Microscope from Leica Microsystems (Wetzlar, Germany). Images were acquired at 63x magnification using an oil immersion objective and Leica LAS-X software. To determine the Piezo1 distribution as well as the channel density, the number of ion channels was collected in 5 µm^2^ regions. Where possible, four regions in the center of the cell and eight regions at the cell edges (periphery) were evaluated per cell. Moreover, background was subtracted from the cellular signal by counting stained structures in 5 µm^2^ regions outside the cell borders. To verify whether a structure was an ion channel, a line scan was placed through the long axis of the corresponding object in ImageJ. A structure was scored as a channel if the associated half-maximal intensity signal extended over an apparent distance of < 4 pixels (< 0.72 µm).

Piezo1^WT^ and Piezo1^GFAP^ knockout PSCs were fixed with 4% paraformaldehyde for 10 minutes at room temperature and then treated with 0.1% Triton X-100 (Swain and Liddle, 2021). Cells were immunostained with a chicken anti-GFAP antibody (ab4674, RRID:AB_304558, Abcam) or rabbit anti-Cre antibody (NB100-56133, RRID:AB_838060, Novus Biological) at 2°C–8°C. Secondary antibodies included DyLight 488–conjugated anti–chicken IgG (703-546-155, RRID: AB_2340376, Jackson ImmunoResearch,), or Cy 3-conjugated anti–rabbit IgG (111-165-003, RRID: AB_2338000, Jackson ImmunoResearch), used for 1 hour at room temperature. All images were captured with a Zeiss Axio observer Z1 with a 20× objective.

### Intracellular Ca^2+^ measurements

For intracellular Ca^2+^ measurements, the ratiometric Ca^2+^ indicator dye Fura-2 was used. The measurements were made using an inverted Axiovert 200 M microscope from Carl Zeiss AG equipped with a pco.edge 5.5 camera (PCO AG, Kelheim, Germany). The images were taken at 40x objective magnification. The VisiChrome High-Speed Polychromator System from Visitron Systems GmbH (Puchheim, Germany) was used as the light source, generating monochromatic light at wavelengths 340 and 380 nm for Fura-2 excitation. The images were taken exclusively in the 4 mm^2^ square, where the presence of the stiffness gradient had been verified using the AFM technique. A gel region was selected in which the Ca^2+^ concentration of the cells was measured over an 8-minute period. For this purpose, recordings were made at 10-second intervals using both excitation wavelengths. First, the glass-bottom dishes were superfused for 2 min with Ca^2+^-free solution (122.5 mM NaCl, 0.8 mM MgCl_2_, 5.4 mM KCl, 10 mM HEPES, 1 mM EGTA),to identify stressed or dying cells, before switching back to HEPES-buffered Ringer’s solution (122.5 mM NaCl, 1.2 mM CaCl_2_, 0.8 mM MgCl_2_, 5.4 mM KCl, 10 mM HEPES, 5.5 mM glucose) for 1 minute. This was followed by a switch to a Ringer’s solution containing 5 µM Yoda1 for 5 minutes. ImageJ 1.52p software was used to evaluate the measurements. First, a pixel-per-pixel background correction was performed, as well as the application of a Gaussian Blur filter with a radius of 200 pixels to reduce background noise. The absolute intensity values of the pixels in both excitation wavelengths were used for the subsequent pixel-by-pixel 340/380 ratio generation. The resulting 32-bit floating-point ratio image was used for all subsequent evaluations. The increase in the F340/F380 ratio served as a measure of the increase in intracellular Ca^2+^ concentration upon Yoda1 administration.

For Piezo1^WT^ and Piezo1^GFAP^ knockout PSCs, cell calcium imaging was performed on quiescent PSCs cultured on the Matrigel-coated plates using Calcium 6-QF dye (Molecular Devices) as previously described (Swain and Liddle, 2021; Swain et al., 2022). Cells were imaged in HBSS buffer with 2 mM Ca^2+^. Imaging was performed using a Zeiss Axio observer Z1 with a 20× objective at room temperature, and the intensity of Calcium 6-QF over time was analyzed using MetaMorph image processing and analysis software (Molecular Devices). Faintly and highly fluorescent-loaded cells were excluded from the analysis. The chemicals used in calcium imaging experiments included Yoda1 (Tocris, cat. No. 5586).

To assess the rate of Ca^2+^-influx into PSCs, we applied the Mn^2+^-quench technique (Fels et al., 2016; Kuntze et al., 2020). Mn^2+^ enters cells with via similar pathways as Ca^2+^. However, unlike Ca^2+^ ions, Mn^2+^ ions quench the fluorescence emission of the Ca^2+^-sensitive dye Fura-2. PSCs were loaded with 3 μM Fura-2 AM for 30 min with DMEM/Ham’s-F12 in the incubator. The experiments were performed at the Ca^2+^ insensitive, isosbestic excitation wavelength of Fura-2 (365 nm), and fluorescence emission was recorded at 510 nm. Images were acquired in 5 s intervals. During measurements, a 5-min control period with Ringer solution was followed by a 5-min perfusion of Mn^2+^-containing Ringer’s (Mn^2+^ Ringer’s) solution. The Mn^2+^ concentration in Mn^2+^ Ringer’s was 200 µM. Data analysis was performed by measuring fluorescence intensities over the whole cell area and correcting it for background fluorescence. For data analysis the intensity was normalized to the values of the control period. Afterwards, regression analysis of the calcium-dependent Fura-2 AM fluorescence intensity over time allowed the determination of the change of fluorescence quenching. The corresponding mean change of the slope (Δm F_365_/t = m2 − m1) was calculated (m1 = slope during control conditions; m2 = initial slope during Mn^2+^ perfusion). Lastly, for easier interpretation, the inverse value of the Mn^2+^ quench was determined which is a surrogate of Ca^2+^ influx.

### Numerical simulations

System (IIIa,IIIb) was numerically solved by a finite volume scheme which is based on the Scharfetter-Gummel flux approximation and is described in more detail in the numerical appendix. The implementation is based on Julia (Bezanson et al., 2017). The simulations in Figure 5 use 16,384 time steps with a step size of 1/5120 on a grid with a spacing of 1/512.

### Statistical analysis

All data sets were tested for normal distribution using the Anderson-Darling and D’Agostino-Pearson tests. For a normally distributed data set, mean and standard error of the mean (SEM) were used to represent the measured values. The t-test was used as a significance test for two comparison groups. In the absence of a normal distribution, the median was mapped, and the 95% confidence interval was mapped as the measure of dispersion. The Mann-Whitney test was used as a hypothesis test for two comparison groups and the Kruskal-Wallis test for multiple comparisons.

All statistical analyses were performed as two-sided tests at a significance level α of 0.05. Dunn’s test was used as a post hoc test in the absence of a normal distribution.

## Supplementary material

Supplementary material contains supplementary Figures S1-S5, and supplementary Table S1-S2, as well as the Numerical appendix.

## Author Contributions

Conceptualization: IB, AlbS, ZP; Methodology: IB, AndS, AK, AngS, SS, SMS; Software: AndS, DI; Investigation: IB, AndS, SS, AK, JMJR, SMS, ZP; Validation: BF, JMJR, RAL, AngS, AlbS, ZP; Formal analysis: AndS, AngS; Visualization: IB, AndS, DI, SMS, ZP; Funding acquisition: RAL, AlbS; Project administration: RAL, AngS, AlbS; Supervision: RAL, AngS, AlbS, ZP; Writing – original draft: IB, AndS, AngS, ZP; Writing – review & editing: AK, BF, JMJR, SMS, RAL, AngS, AlbS, ZP

## Supporting information

Numerics appendix

## Acknowledgments

We would like to thank Prof. Timo Betz, Prof. Hermann Schillers and Dr. Ivan Liashkovich for their helpful insights on hydrogels and atomic force microscopy. IB and AK were supported by MedK of the Medical Faculty of the University of Münster. AS was funded by IZKF Münster (Schw2-020-18), Deutsche Forschungsgemeinschaft (SCHW407/22-1) and the EU (Horizon 2020 Marie Skłodowska-Curie grant pHioniC, No 813834). We acknowledge support from National Institutes of Health award R01DK124474 (RAL). AndS and AngS were supported by the DFG (German-Research-Foundation) under Germany’s Excellence Strategy EXC 2044-390685587, Mathematics Münster: Dynamics – Geometry – Structure.

## References

Acerbi, I., L. Cassereau, I. Dean, Q. Shi, A. Au, C. Park, Y.Y. Chen, J. Liphardt, E.S. Hwang, and V.M. Weaver. 2015. Human breast cancer invasion and aggression correlates with ECM stiffening and immune cell infiltration. Integrative Biology. 7. doi:10.1039/c5ib00040h.

Allena, R., M. Scianna, and L. Preziosi. 2016. A Cellular Potts Model of single cell migration in presence of durotaxis. Math Biosci. 275:57–70. doi:10.1016/J.MBS.2016.02.011.

Apte, M. V., Z. Xu, S. Pothula, D. Goldstein, R.C. Pirola, and J.S. Wilson. 2015. Pancreatic cancer: The microenvironment needs attention too! Pancreatology. 15. doi:10.1016/j.pan.2015.02.013.

Bae, C., F. Sachs, and P.A. Gottlieb. 2011. The mechanosensitive ion channel Piezo1 is inhibited by the peptide GsMTx4. Biochemistry. 50:6295–6300. doi:10.1021/bi200770q.

Bera, K., A. Kiepas, I. Godet, Y. Li, P. Mehta, B. Ifemembi, C.D. Paul, A. Sen, S.A. Serra, K. Stoletov, J. Tao, G. Shatkin, S.J. Lee, Y. Zhang, A. Boen, P. Mistriotis, D.M. Gilkes, J.D. Lewis, C.M. Fan, A.P. Feinberg, M.A. Valverde, S.X. Sun, and K. Konstantopoulos. 2022. Extracellular fluid viscosity enhances cell migration and cancer dissemination. Nature 2022 611:7935. 611:365–373. doi:10.1038/s41586-022-05394-6.

Bezanson, J., A. Edelman, S. Karpinski, and V.B. Shah. 2017. Julia: A Fresh approach to numerical computing. 10.1137/141000671. 59:65–98. doi:10.1137/141000671.

Cook, S.M., T.E. Schäffer, K.M. Chynoweth, M. Wigton, R.W. Simmonds, and K.M. Lang. 2006. Practical implementation of dynamic methods for measuring atomic force microscope cantilever spring constants. Nanotechnology. 17. doi:10.1088/0957-4484/17/9/010.

Coste, B., J. Mathur, M. Schmidt, T.J. Earley, S. Ranade, M.J. Petrus, A.E. Dubin, and A. Patapoutian. 2010. Piezo1 and Piezo2 are essential components of distinct mechanically activated cation channels. Science (1979). 330. doi:10.1126/science.1193270.

Dietrich, A., H. Kalwa, U. Storch, M. Mederos, Y Schnitzler, B. Salanova, O. Pinkenburg, G. Dubrovska, K. Essin, M. Gollasch, L. Birnbaumer, and T. Gudermann. 2007. Pressure-induced and store-operated cation influx in vascular smooth muscle cells is independent of TRPC1. Pflugers Arch. 455. doi:10.1007/s00424-007-0314-3.

DuChez, B.J., A.D. Doyle, E.K. Dimitriadis, and K.M. Yamada. 2019. Durotaxis by human cancer cells. Biophys J. 116. doi:10.1016/j.bpj.2019.01.009.

Ellefsen, K.L., J.R. Holt, A.C. Chang, J.L. Nourse, J. Arulmoli, A.H. Mekhdjian, H. Abuwarda, F. Tombola, L.A. Flanagan, A.R. Dunn, I. Parker, and M.M. Pathak. 2019. Myosin-II mediated traction forces evoke localized Piezo1-dependent Ca^2+^ flickers. Commun Biol. 2. doi:10.1038/s42003-019-0514-3.

Espina, J.A., C.L. Marchant, and E.H. Barriga. 2022. Durotaxis: the mechanical control of directed cell migration. FEBS J. 289:2736–2754. doi:10.1111/FEBS.15862.

Evans, E.B., S.W. Brady, A. Tripathi, and D. Hoffman-Kim. 2018. Schwann cell durotaxis can be guided by physiologically relevant stiffness gradients. Biomater Res. 22. doi:10.1186/s40824-018-0124-z.

Everaerts, W., X. Zhen, D. Ghosh, J. Vriens, T. Gevaert, J.P. Gilbert, N.J. Hayward, C.R. McNamara, F. Xue, M.M. Moran, T. Strassmaier, E. Uykal, G. Owsianik, R. Vennekens, D. De Ridder, B. Nilius, C.M. Fanger, and T. Voets. 2010. Inhibition of the cation channel TRPV4 improves bladder function in mice and rats with cyclophosphamide-induced cystitis. Proc Natl Acad Sci U S A. 107:19084–19089. doi:10.1073/PNAS.1005333107/SUPPL_FILE/PNAS.201005333SI.PDF.

Fabian, A., J. Bertrand, O. Lindemann, T. Pap, and A. Schwab. 2012. Transient receptor potential canonical channel 1 impacts on mechanosignaling during cell migration. Pflugers Arch. 464:623– 630. doi:10.1007/S00424-012-1169-9.

Fabian, A., T. Fortmann, P. Dieterich, C. Riethmüller, P. Schön, S. Mally, B. Nilius, and A. Schwab. 2008. TRPC1 channels regulate directionality of migrating cells. Pflugers Arch. 457. doi:10.1007/s00424-008-0515-4.

Fels, B., N. Nielsen, and A. Schwab. 2016. Role of TRPC1 channels in pressure-mediated activation of murine pancreatic stellate cells. Eur Biophys J. 45:657–670. doi:10.1007/S00249-016-1176-4.

Goswami, C., J. Kuhn, P.A. Heppenstall, and T. Hucho. 2010. Importance of non-selective cation channel TRPV4 interaction with cytoskeleton and their reciprocal regulations in cultured cells. PLoS One. 5. doi:10.1371/journal.pone.0011654.

Gottlieb, P., J. Folgering, R. Maroto, A. Raso, T.G. Wood, A. Kurosky, C. Bowman, D. Bichet, A. Patel, F. Sachs, B. Martinac, O.P. Hamill, and E. Honoré. 2008. Revisiting TRPC1 and TRPC6 mechanosensitivity. Pflugers Arch. 455. doi:10.1007/s00424-007-0359-3.

Holt, J.R., W.Z. Zeng, E.L. Evans, S.H. Woo, S. Ma, H. Abuwarda, M. Loud, A. Patapoutian, and M.M. Pathak. 2021. Spatiotemporal dynamics of piezo1 localization controls keratinocyte migration during wound healing. Elife. 10. doi:10.7554/eLife.65415.

Janmey, P.A., D.A. Fletcher, and C.A. Reinhart-King. 2020. Stiffness sensing by cells. Physiol Rev. 100. doi:10.1152/physrev.00013.2019.

Kobayashi, T., and M. Sokabe. 2010. Sensing substrate rigidity by mechanosensitive ion channels with stress fibers and focal adhesions. Curr Opin Cell Biol. 22. doi:10.1016/j.ceb.2010.08.023.

Kuntze, A., O. Goetsch, B. Fels, K. Najder, A. Unger, M. Wilhelmi, S. Sargin, S. Schimmelpfennig, I. Neumann, A. Schwab, and Z. Pethő. 2020. Protonation of piezo1 impairs cell-matrix interactions of pancreatic stellate cells. Front Physiol. 11:89. doi:10.3389/FPHYS.2020.00089/BIBTEX.

Lachowski, D., E. Cortes, D. Pink, A. Chronopoulos, S.A. Karim, J.P. Morton, and A.E. del R. Hernández. 2017. substrate rigidity controls activation and durotaxis in pancreatic stellate cells. Sci Rep. 7. doi:10.1038/S41598-017-02689-X.

Lachowski, D., E. Cortes, B. Robinson, A. Rice, K. Rombouts, and A.E. Del Río Hernández. 2018. FAK controls the mechanical activation of YAP, a transcriptional regulator required for durotaxis. FASEB J. 32:1099–1107. doi:10.1096/FJ.201700721R.

Lo, C.M., H.B. Wang, M. Dembo, and Y.L. Wang. 2000. Cell movement is guided by the rigidity of the substrate. Biophys J. 79. doi:10.1016/S0006-3495(00)76279-5.

Malik, A.A., and P. Gerlee. 2019. Mathematical modelling of cell migration: stiffness dependent jump rates result in durotaxis. J Math Biol. 78:2289–2315. doi:10.1007/S00285-019-01344-5.

Maroto, R., A. Raso, T.G. Wood, A. Kurosky, B. Martinac, and O.P. Hamill. 2005. TRPC1 forms the stretch-activated cation channel in vertebrate cells. Nat Cell Biol. 7. doi:10.1038/ncb1218.

Matthews, B.D., C.K. Thodeti, J.D. Tytell, A. Mammoto, D.R. Overby, and D.E. Ingber. 2010. Ultra-rapid activation of TRPV4 ion channels by mechanical forces applied to cell surface β1 integrins. Integrative Biology. 2. doi:10.1039/c0ib00034e.

Nielsen, N., K. Kondratska, T. Ruck, B. Hild, I. Kovalenko, S. Schimmelpfennig, J. Welzig, S. Sargin, O. Lindemann, S. Christian, S.G. Meuth, N. Prevarskaya, and A. Schwab. 2017. TRPC6 channels modulate the response of pancreatic stellate cells to hypoxia. Pflugers Arch. 469:1567–1577. doi:10.1007/S00424-017-2057-0.

Nikolaev, Y.A., C.D. Cox, P. Ridone, P.R. Rohde, J.F. Cordero-Morales, V. Vásquez, D.R. Laver, and B. Martinac. 2019. Mammalian TRP ion channels are insensitive to membrane stretch. J Cell Sci. 132. doi:10.1242/jcs.238360.

Nourse, J.L., and M.M. Pathak. 2017. How cells channel their stress: Interplay between Piezo1 and the cytoskeleton. Semin Cell Dev Biol. 71. doi:10.1016/j.semcdb.2017.06.018.

Othmer, H.G., and A. Stevens. 2006. Aggregation, blowup, and collapse: The ABC’s of taxis in reinforced random walks. 10.1137/S0036139995288976. 57:1044–1081. doi:10.1137/S0036139995288976.

Pethő, Z., K. Najder, S. Beel, B. Fels, I. Neumann, S. Schimmelpfennig, S. Sargin, M. Wolters, K. Grantins, E. Wardelmann, M. Mitkovski, A. Oeckinghaus, and A. Schwab. 2023. Acid-base homeostasis orchestrated by NHE1 defines pancreatic stellate cell phenotype in pancreatic cancer. JCI Insight. doi:10.1172/JCI.INSIGHT.170928.

Pethő, Z., K. Najder, E. Bulk, and A. Schwab. 2019. Mechanosensitive ion channels push cancer progression. Cell Calcium. 80:79–90. doi:10.1016/j.ceca.2019.03.007.

Piersma, B., M.K. Hayward, and V.M. Weaver. 2020. Fibrosis and cancer: A strained relationship. Biochim Biophys Acta Rev Cancer. 1873. doi:10.1016/j.bbcan.2020.188356.

Pothula, S.P., R.C. Pirola, J.S. Wilson, and M. V. Apte. 2020. Pancreatic stellate cells: Aiding and abetting pancreatic cancer progression. Pancreatology. 20:409–418. doi:10.1016/J.PAN.2020.01.003.

Radoslavova, S., B. Fels, Z. Pethö, M. Gruner, T. Ruck, S.G. Meuth, A. Folcher, N. Prevarskaya, A. Schwab, and H. Ouadid-Ahidouch. 2022. TRPC1 channels regulate the activation of pancreatic stellate cells through ERK1/2 and SMAD2 pathways and perpetuate their pressure-mediated activation. Cell Calcium. 106. doi:10.1016/J.CECA.2022.102621.

Rheinlaender, J., A. Dimitracopoulos, B. Wallmeyer, N.M. Kronenberg, K.J. Chalut, M.C. Gather, T. Betz, G. Charras, and K. Franze. 2020. Cortical cell stiffness is independent of substrate mechanics. Nature Materials 2020 19:9. 19:1019–1025. doi:10.1038/s41563-020-0684-x.

Romac, J.M.J., R.A. Shahid, S.M. Swain, S.R. Vigna, and R.A. Liddle. 2018. Piezo1 is a mechanically activated ion channel and mediates pressure induced pancreatitis. Nat Commun. 9. doi:10.1038/s41467-018-04194-9.

Rubiano, A., D. Delitto, S. Han, M. Gerber, C. Galitz, J. Trevino, R.M. Thomas, S.J. Hughes, and C.S. Simmons. 2018. Viscoelastic properties of human pancreatic tumors and in vitro constructs to mimic mechanical properties. Acta Biomater. 67. doi:10.1016/j.actbio.2017.11.037.

Schwab, A., B. Schuricht, P. Seeger, J. Reinhardt, and P.C. Dartsch. 1999. Migration of transformed renal epithelial cells is regulated by K^+^ channel modulation of actin cytoskeleton and cell volume. Pflugers Arch. 438. doi:10.1007/s004240050917.

Sheth, S., E. Jain, A. Karadaghy, S. Syed, H. Stevenson, and S.P. Zustiak. 2017. UV dose governs UV-polymerized polyacrylamide hydrogel modulus. Int J Polym Sci. 2017. doi:10.1155/2017/5147482.

Shi, Y., L. Cang, X. Zhang, X. Cai, X. Wang, R. Ji, M. Wang, and Y. Hong. 2018. The use of magnetic resonance elastography in differentiating autoimmune pancreatitis from pancreatic ductal adenocarcinoma: A preliminary study. Eur J Radiol. 108. doi:10.1016/j.ejrad.2018.09.001.

Stock, C., B. Gassner, C.R. Hauck, H. Arnold, S. Mally, J.A. Eble, P. Dieterich, and A. Schwab. 2005. Migration of human melanoma cells depends on extracellular pH and Na^+^/H^+^ exchange. J Physiol. 567:225. doi:10.1113/JPHYSIOL.2005.088344.

Storck, H., B. Hild, S. Schimmelpfennig, S. Sargin, N. Nielsen, A. Zaccagnino, T. Budde, I. Novak, H. Kalthoff, and A. Schwab. 2017. Ion channels in control of pancreatic stellate cell migration. Oncotarget. 8:769–784. doi:10.18632/ONCOTARGET.13647.

Sun, B. 2021. The mechanics of fibrillar collagen extracellular matrix. Cell Rep Phys Sci. 2. doi:10.1016/j.xcrp.2021.100515.

Swain, S.M., and R.A. Liddle. 2021. Piezo1 acts upstream of TRPV4 to induce pathological changes in endothelial cells due to shear stress. Journal of Biological Chemistry. 296. doi:10.1074/jbc.RA120.015059.

Swain, S.M., J.M.J. Romac, R.A. Shahid, S.J. Pandol, W. Liedtke, S.R. Vigna, and R.A. Liddle. 2020. TRPV4 channel opening mediates pressure-induced pancreatitis initiated by Piezo1 activation. Journal of Clinical Investigation. 130. doi:10.1172/JCI134111.

Swain, S.M., J.M.J. Romac, S.R. Vigna, and R.A. Liddle. 2022. Piezo1-mediated stellate cell activation causes pressure-induced pancreatic fibrosis in mice. JCI Insight. 7. doi:10.1172/jci.insight.158288.

Syeda, R., M.N. Florendo, C.D. Cox, J.M. Kefauver, J.S. Santos, B. Martinac, and A. Patapoutian. 2016. Piezo1 channels are inherently mechanosensitive. Cell Rep. 17:1739–1746. doi:10.1016/J.CELREP.2016.10.033.

Syeda, R., J. Xu, A.E. Dubin, B. Coste, J. Mathur, T. Huynh, J. Matzen, J. Lao, D.C. Tully, I.H. Engels, H. Michael Petrassi, A.M. Schumacher, M. Montal, M. Bandell, and A. Patapoutian. 2015. Chemical activation of the mechanotransduction channel Piezo1. Elife. 4. doi:10.7554/eLife.07369.

Thorneloe, K.S., A.C. Sulpizio, Z. Lin, D.J. Figueroa, A.K. Clouse, G.P. McCafferty, T.P. Chendrimada, E.S.R. Lashinger, E. Gordon, L. Evans, B.A. Misajet, D.J. DeMarini, J.H. Nation, L.N. Casillas, R.W. Marquis, B.J. Votta, S.A. Sheardown, X. Xu, D.P. Brooks, N.J. Laping, and T.D. Westfall. 2008. GSK1016790A, a novel and potent transient receptor potential vanilloid 4 channel agonist induces urinary bladder contraction and hyperactivity: Part I [S]. Journal of Pharmacology and Experimental Therapeutics. 326. doi:10.1124/jpet.108.139295.

Tse, J.R., and A.J. Engler. 2010. Preparation of hydrogel substrates with tunable mechanical properties. Curr Protoc Cell Biol. doi:10.1002/0471143030.cb1016s47.

Waschk, D.E.J., A. Fabian, T. Budde, and A. Schwab. 2011. Dual-color quantum dot detection of a heterotetrameric potassium channel (hK ca3.1). Am J Physiol Cell Physiol. 300. doi:10.1152/ajpcell.00053.2010.

Wei, C., X. Wang, M. Chen, K. Ouyang, L.S. Song, and H. Cheng. 2009. Calcium flickers steer cell migration. Nature. 457. doi:10.1038/nature07577.

