## Supporting Figures and Table for "Piezo1-induced durotaxis of pancreatic stellate cells depends on TRPC1 and TRPV4 channels"

**This PDF file includes:**

Figures S1 to S6
Table S1
Numerics

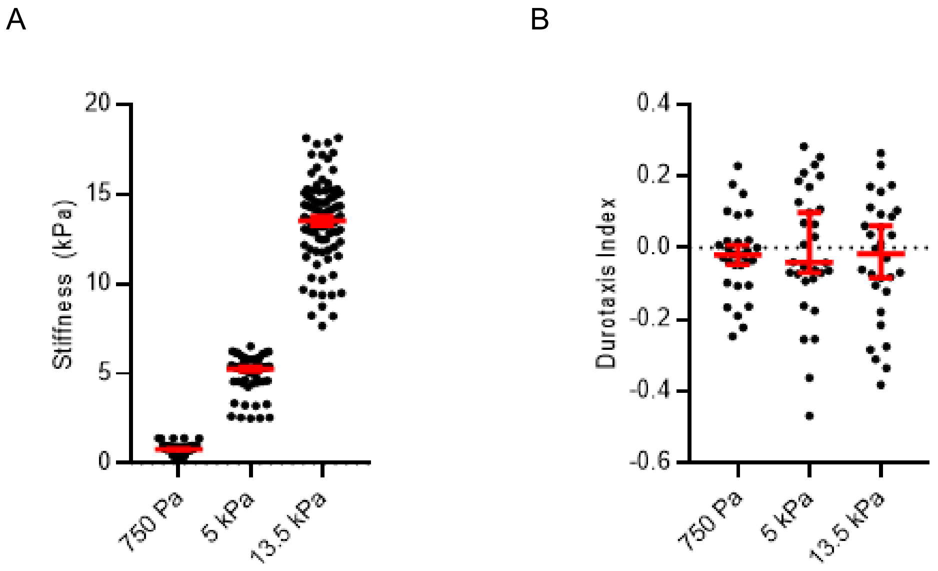


**Fig. S1. PSCs on hydrogels with constant stiffness migrate non-directionally**

**A)** Scatter plot shows homogeneous gel stiffnesses measured from n≥10 points of N=5 gels. **B)** Scatter plot indicates durotaxis indices of PSCs derived from the trajectories in Fig. 1B. (n=30 PSCs in N=3 independent experiments). Data in (A) is mean ± SEM and in (B) median ± 95% CI. Statistical test in (B) is one-sample t-test.

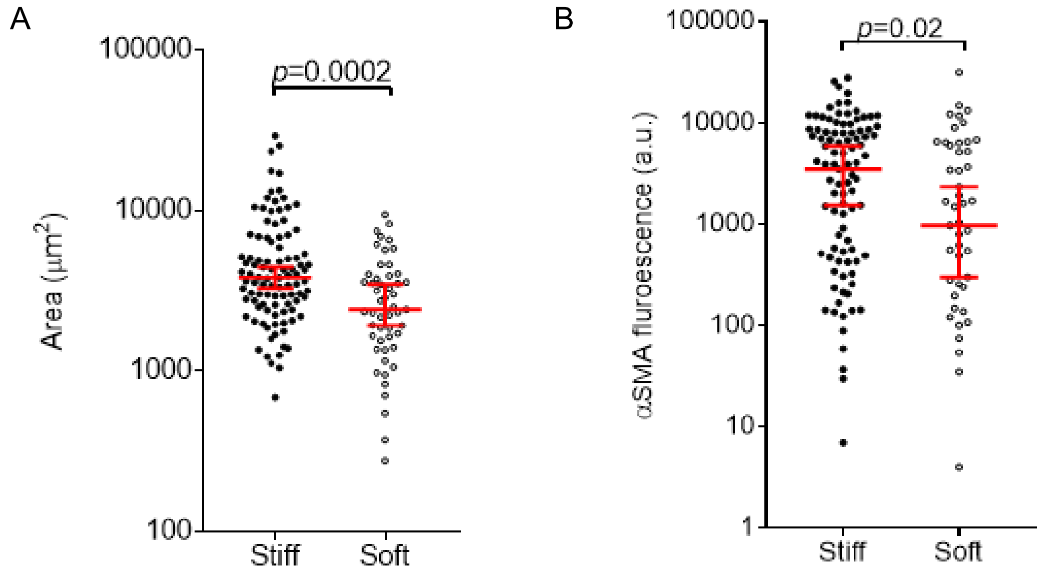


**Fig. S2. PSC immunostaining reveals higher cell area and αSMA intensity on a stiffer substrate**

**A)** Scatter plot shows cellular areas derived from αSMA-stained PSCs depicted in Fig. 2E. **B)** Scatter plot of αSMA fluorescence intensity of PSCs on the stiff and soft parts of the gradient hydrogel. n cells measured / N experiments ≥ 51/4. Data in (A) and (B) are median ± 95% CI. Statistical test are Mann-Whitney U-tests.

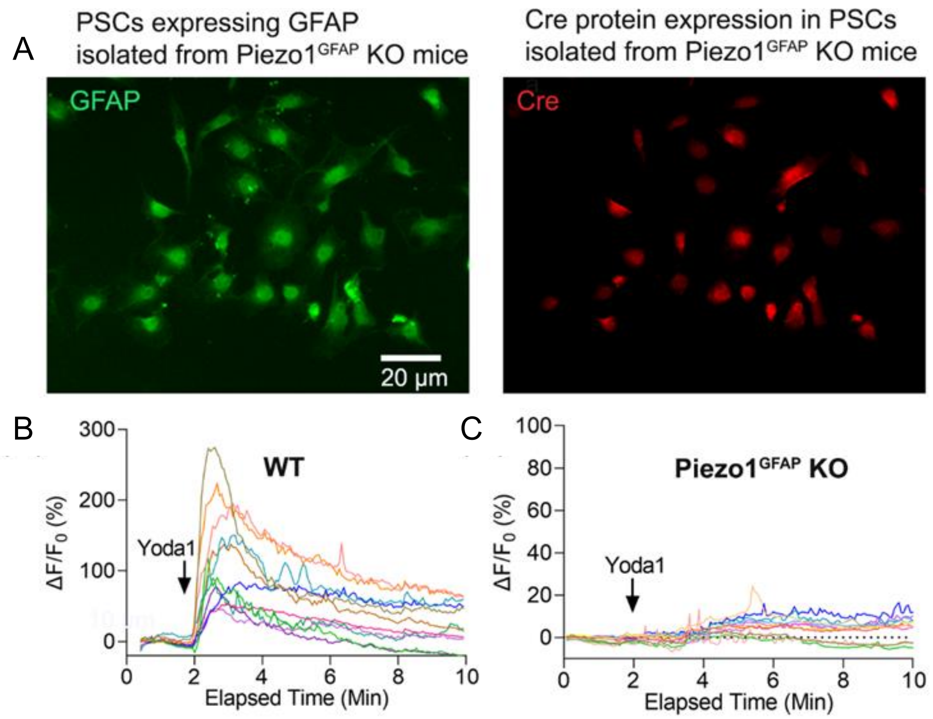


**Fig. S3. Piezo1 deletion in GFAP expressing mouse PSCs**

**A)** Images showing PSCs isolated from mouse line B6. Cg-Tg (GFAP- cre/ERT2; *Piezo1$^{fl/f}$)* after tamoxifen injection (referred to as Piezo1$^{GFAP}$ KO mice) cultured for 3 days in a Matrigel-coated plate as described previously (1), expressed both stellate cell marker GFAP (green) and Cre protein (red). Scale bar: 20 µm. **B)** and **C)** Piezo1 agonist Yoda1 (5 µM) induces an elevation of the intracellular $Ca^{2+}$ concentration in PSCs isolated from wild type but not from Piezo1$^{GFAP}$ KO mice, confirming that successful deletion of Piezo1 in GFAP expressing PSCs, n=12 cells. Each trace represents Yoda1-induced single cell relative fluorescence intensity ($\Delta F/F_0$) of the calcium 6-QF dye over time. $\Delta F$ is the change in fluorescence intensity ($F\text{-}F_0$), and $F_0$ is the basal fluorescence intensity before Yoda1 application.

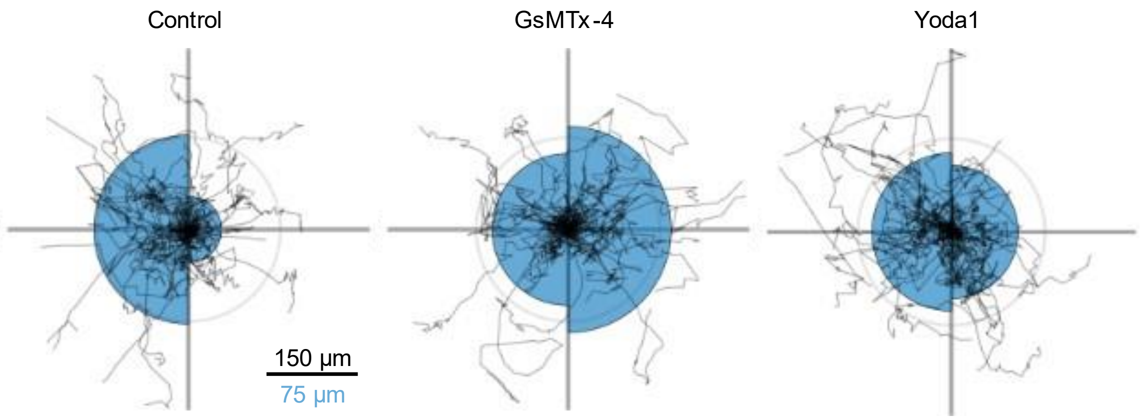


**Fig. S4. Durotaxis polar plots of PSCs with pharmacological Piezo1 modulation**

The durotaxis plot data aims to support Fig. 3C. Durotaxis polar plots depict individual PSC trajectories over 24 h (black lines). The radius of the blue half circles is proportional to the mean cellular displacement into the directions 0° and 180°, respectively. Radial lines indicate 0°, 90°, 180°, 270°.

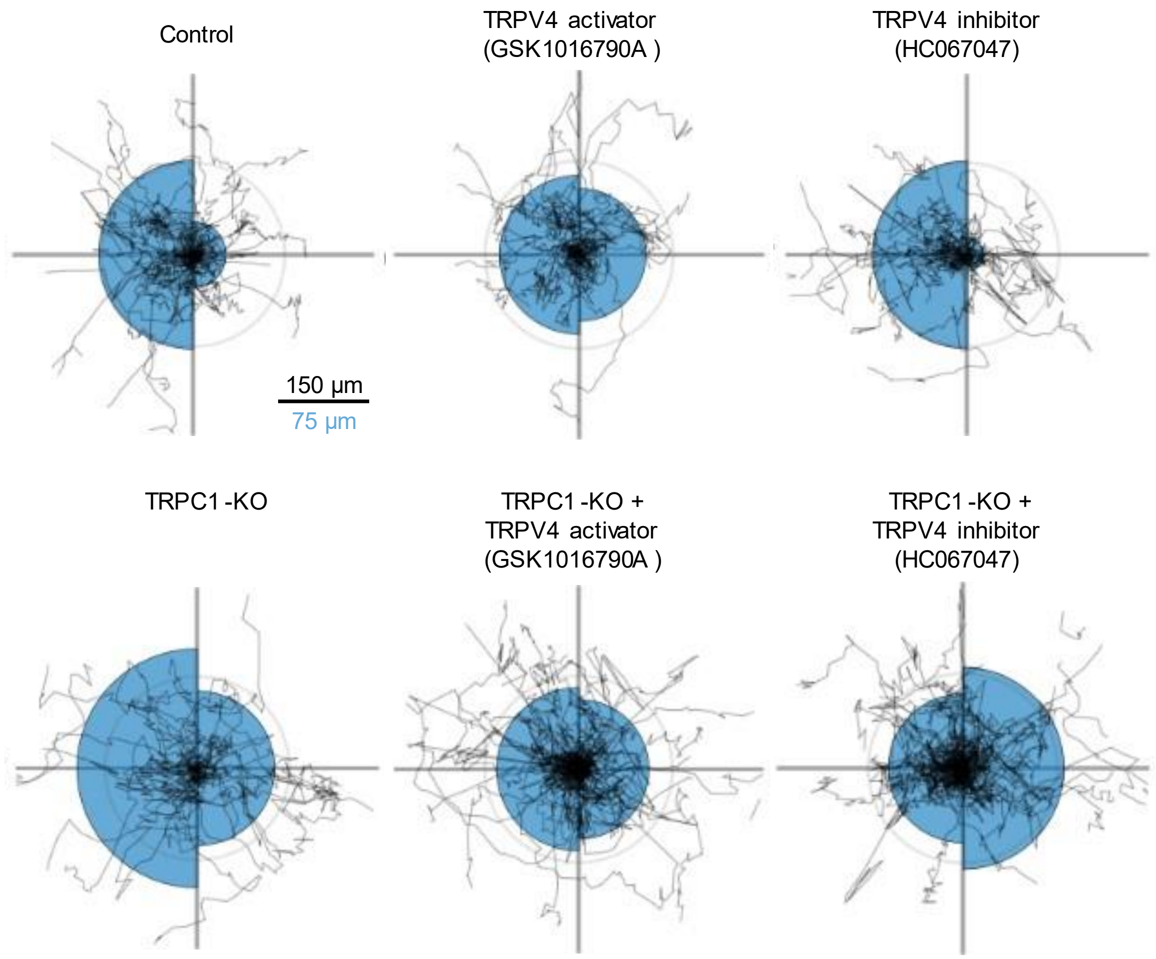


**Fig. S5. Durotaxis polar plots of PSCs with TRPV4 and TRPC1 modulation**

The durotaxis plot data aims to support Fig. 5A. Durotaxis polar plots depict individual PSC trajectories over 24 h (black lines). The radius of the blue half circles is proportional to the mean cellular displacement into the directions 0° and 180°, respectively. Radial lines indicate 0°, 90°, 180°, 270°.

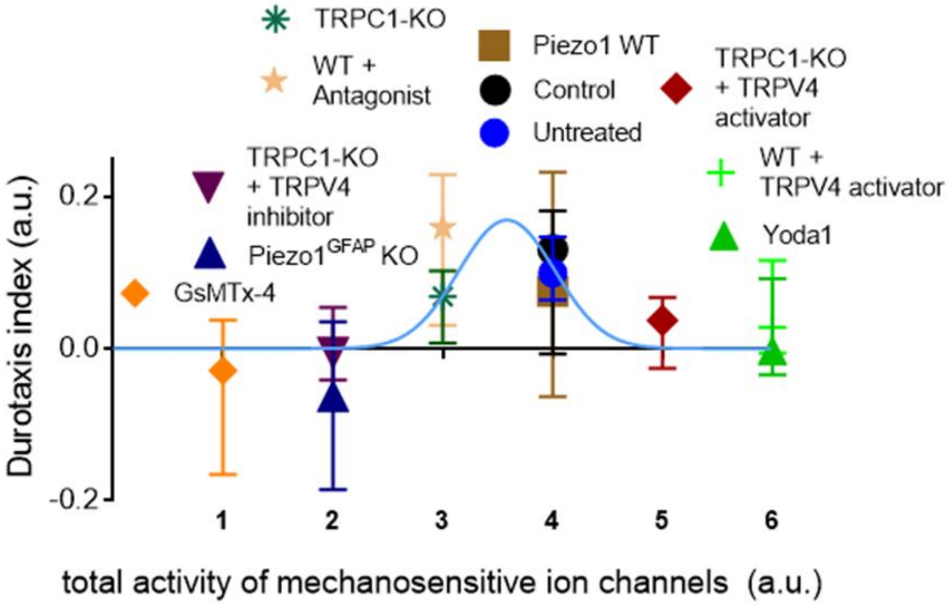


**Fig. S6. Durotaxis requires intermediate activity of mechanosensitive ion channels**

The depicted data shows details regarding Fig. 5C. Scatter plot shows the durotaxis index as a function of the total ion channel activity, derived from Fig. 5A and B. Inhibitor and activator refer to TRPV4 modulation with HC-0606006 and GSK101489, respectively. The blue line corresponds to a gaussian curve fit from Fig. 5c, indicating the bell-shaped relationship between channel activity and durotaxis. Data points are median ± 95% CI.

### Supporting Tables

**Table S1.** Acrylamide mixture used for hydrogels with constant stiffness

| Acrylamide (%) | Bisacrylamide (%) | Hydroxyacryl-amide (%) | Volume Acrylamide mix (µl) | Volume PBS (µl) | Stiffness |
|---|---|---|---|---|---|
| 2,5 | 0,07 | 0,8 | 53 | 447 | 750 Pa |
| 3,54 | 0,1 | 1,15 | 75 | 425 | 5 kPa |
| 7,08 | 0,2 | 2,3 | 150 | 350 | 13.5 kPa |
