## Supplementary material for "Piezo1-induced durotaxis of pancreatic stellate cells depends on TRPC1 and TRPV4 channels": Numerics appendix

### Appendix - Numerical Simulation of (IIIa)–(IIIb)

#### 1 Finite-volume discretization

The numerical simulation of the system (IIIa)–(IIIb) given by

$$\frac{\partial}{\partial t}\rho = \nabla \cdot \left( \frac{\rho}{\rho+\eta} \nabla(\rho+\eta) - \rho q \left( \frac{\rho}{\rho+\eta} \right) \nabla E \right) + R(\rho,\eta) \tag{IIIa}$$

$$\frac{\partial}{\partial t}\eta = \nabla \cdot \left( \frac{\eta}{\rho+\eta} \nabla(\rho+\eta) - \eta q \left( \frac{\rho}{\rho+\eta} \right) \nabla E \right) - R(\rho,\eta), \tag{IIIb}$$

is based on a finite volume scheme described below. Here $\rho = \rho(x,t)$ , $\eta = \eta(x,t)$ denote the densities of the open and closed channels on the cell respectively. $E$ denotes the elastic modulus of the substratum, i.e. it is larger for a stiffer material. The function $q$ accounts for the cell's strength of sensing the gradient of $E$ due to the relative number of open versus closed channels. $R(\rho,\eta)$ describes the combined opening and closing of the channels. In our mathematical model we defined $R(\rho,\eta) = \alpha E \cdot \eta - \beta\rho$.

The proposed numerical scheme to solve the above system is based on the Scharfetter–Gummel flux approximation originating from (13), where the authors construct a numerical scheme for a system modelling semiconductor devices. Their objective was to develop a robust scheme for discontinuities or rapid variations in the potential. Independently, the same type of flux was introduced in (10) for finite-difference schemes. The Scharfetter–Gummel scheme became one of the preferred finite-volume scheme for drift-diffusion equations. While the original scheme deals with the spatially one-dimensional problem, it has been generalized to higher dimensions (8; 9) and the flux discretization is the basis for numerous other generalizations, e.g. for equations with nonlinear diffusion (11; 3; 5) and to systems with source terms (4; 16). In the context of chemotaxis and aggregation models the Schafetter–Gummel flux approximation was recently used in (17; 12; 1; 15; 14) see also the review (2).

First we generalize our problem to

$$\partial_t \rho = \nabla \cdot \left( \frac{\rho}{\rho+\eta} J[\rho+\eta] \right) + R(\rho,\eta) \tag{IVa}$$

$$\partial_t \eta = \nabla \cdot \left( \frac{\eta}{\rho+\eta} J[\rho+\eta] \right) - R(\rho,\eta), \tag{IVb}$$

which is in flux-form. The we set $J[u] = \nabla u + uW$ for some given (possibly time-dependent) vectorfield $W$. This will later be defined as a numerical approximation of $-q(p(x,t))\nabla E(x)$. Hence, the above driving flux $J$ depends only on the sum $u = \rho + \eta$ and space and time.

The notation for finite volume schemes is as follows (see e.g. (6)). Consider a polyhedral tesselation $\mathcal{T}$ of the domain $\Omega$ with volumes $K \in \mathcal{T}$ and centers $x_K$. Neighboring cells are denoted by $L \sim K$ with common face $K|L := \overline{K} \cap \overline{L}$.
*Note: here cells are a technical expression for so-called numerical cells in finite volume schemes, which have nothing to do with the biological cells we are considering in this paper.*
The distance between cell centers is denoted with $d_{KL} := |x_k - x_L|$.
The transmission coefficient is $\tau_{KL} = \frac{|K|L|}{d_{KL}} = \tau_{LK}$.
We use $|K|$ to denote the $d$-dimensional volume and by slight abuse of notation we denote by $|K|L|$ the $d-1$-dimensional surface area.
We use $\rho_K, \eta_K$ for the cell-averages of densities and set $u_K = \rho_K + \eta_K$.
$p_K = \frac{\rho_K}{u_K}$ and $q_K = \frac{\eta_K}{u_K}$ denote the relative weights of the two densities.
For $\tau \in [0,1]$, $\rho^{n+\tau}$ denotes the linear-interpolation between $\rho^n$ and $\rho^{n+1}$.
This allows to define the full range of explicit to implicit schemes at the same time. Given some flux-approximation $J_{KL}[u^{n+\tau}]$ which will be specified later, we introduce the scheme

$$\frac{\rho_K^{n+1} - \rho_K^n}{\delta} = \sum_{L\sim K} \tau_{KL}\big(p_L^{n+\tau} J_{KL}[u^{n+\tau}]_- - p_K^{n+\tau} J_{KL}[u^{n+\tau}]_+\big) + R_K(\rho_K^{n+\tau}, \eta_K^{n+\tau}) \tag{Va}$$

$$\frac{\eta_K^{n+1} - \eta_K^n}{\delta} = \sum_{L\sim K} \tau_{KL}\big(q_L^{n+\tau} J_{KL}[u^{n+\tau}]_- - q_K^{n+\tau} J_{KL}[u^{n+\tau}]_+\big) - R_K(\rho_K^{n+\tau}, \eta_K^{n+\tau}), \tag{Vb}$$

where $(a)_+ = \max\{a, 0\}$ and $(a)_- = (-a)_+$ denote the positive and negative part, respectively. Our reaction term is explicitly given by

$$R_K(\rho, \eta) = \alpha(E_K)\eta - \beta\rho,$$

with $E_K$ being the cell-average of the elastic module $E$.

It is left to find approximations for the normal fluxes $J_{KL}^{n+\tau}$ across boundary faces. If we assume that some discretization $W_{KL}^{n+\tau}$ of $W$ (depending in our case on $p_K^{n+\tau}$) is given, then the normal component of the flux $J[u]$ in (IVa)–(IVb) can be approximated by the cell problem (see e.g. (13; 11; 7; 14))

$$J_{KL}^{n+\tau} = \partial_x u + u W_{KL}^{n+\tau},$$

where $u$ is the unknown along the line segment $[x_k, x_L] := \{(1-s)x_k + s x_L : s \in [0,1]\}$ with boundary conditions $u(x_k) = u_K^{n+\tau}$ and $u(x_L) = u_L^{n+\tau}$.

This is the classical Scharfetter-Gummel interpolation, and by setting $d_{KL} = |x_K - x_L|$ we obtain the flux $J = J[u_K^{n+\tau}, u_L^{n+\tau}; W_{KL}^{n+\tau}/d_{KL}]$ given by

$$J_{\mathrm{SG}}[a, b; W] = \begin{cases} W \frac{ae^{W/2} - be^{-W/2}}{e^{W/2} - e^{-W/2}}, & W \neq 0; \\ (a - b), & W = 0. \end{cases} \tag{VI}$$

Hence, we can close the scheme by setting

$$J_{KL}[u^{n+\tau}] := J_{\mathrm{SG}}[u_K^{n+\tau}, u_L^{n+\tau}; W_{KL}^{n+\tau}/d_{KL}] \tag{VII}$$

with $W_{KL}^{n+\tau} := W_{KL}[p_K^{n+\tau}]$ given by

$$W_{KL}[p^{n+\tau}] := q(p_K^{n+\tau})(E_L - E_K)_+ - q(p_L^{n+\tau})(E_L - E_K)_- \tag{VIII}$$

### 2 Simulation

For the simulation in Figure 5, we implemented an explicit version ($\tau = 0$) of the scheme (Va)–(Vb) on the interval $[0, 1]$ with a uniform tesselation. That is for given $h = 1/N$, we set $\mathcal{T}_h = \{[0, kh] : k = 1, \ldots, N\}$ and $x_K = (k + 1/2)h$ for $k = 0, \ldots, N - 1$. We have the two discrete continuity equations

$$\frac{\rho_K^{n+1} - \rho_K^n}{\delta} = \sum_{L \sim K} \tau_{KL}(p_L^n J_{KL}[u^n]_- - p_K^n J_{KL}[u^n]_+) + R_K(\rho_K^n, \eta_K^n)$$
$$\frac{\eta_K^{n+1} - \eta_K^n}{\delta} = \sum_{L \sim K} \tau_{KL}(q_L^n J_{KL}[u^n]_- - q_K^n J_{KL}[u^n]_+) - R_K(\rho_K^n, \eta_K^n),$$

where we impose no-flux boundary condition, that is $J_{0,1} = 0 = J_{N,N+1}$ and the sum over $L$ $K$ consists of the two addends $K - 1$ and $k + 1$. Hereby in one dimension $\tau_{KL} = h^{-1}$ and

$$p_K^n = \frac{\rho_K^n}{\rho_K^n + \eta_K^n} \quad \text{and} \quad q_K^n = \frac{\eta_K^n}{\rho_K^n + \eta_K^n}.$$

For the implementation, we choose

$$R_K(\rho, \eta) = \alpha E_K \eta - \beta \rho \qquad \text{with} \qquad E_K = E(x_K).$$

For the simulation we use an elasticity profile on $[0, 1]$ resembling the elasticity module of the experimental made substrate given by

$$E(x) = 2(E_{\mathrm{hard}} - E_{\mathrm{soft}})B(x - 1/2, s) + E_{\mathrm{hard}}$$

where $B(y, s) = y/(\exp(-s\,y) - 1)$. In the simulations in Figure 5, $E_{\mathrm{hard}} = 0.1$, $E_{\mathrm{soft}} = 0.01$ and the scale parameter $s = 30$. We use the Scharfetter-Gummel flux interpolation (VII) in explicit form

$$J_{KL}[u^n] := J_{\mathrm{SG}}[u_K^n, u_L^n; W_{KL}^n/d_{KL}]$$

with $J_{\mathrm{SG}}$ given in (VI) and upwind discretization of the chemical potential given in (VIII) by

$$W_{KL}[p^n] := q(p_K^n)(E_L - E_K)_+ - q(p_L^n)(E_L - E_K)_-$$

Hereby, the discrete mobility is $q_K^n = q(p_K^n)$ with the continuous mobility function given in terms of a bell-shaped function with compact support by

$$q(x) = \nu(3(x - 0.5)) + 0.1 \qquad \text{with} \qquad \nu(x) = \begin{cases} \exp\Big(-\frac{x^2}{1-x^2}\Big), & x \in (-1,1); \\ 0, & |x| \geq 1. \end{cases}$$

### References


[1] L. Almeida, , F. Bubba, B. Perthame, and C. Pouchol. Energy and implicit discretization of the Fokker-Planck and Keller-Segel type equations. *Networks & Heterogeneous Media*, 14(1):23–41, 2019.

[2] G. Bärwolff and D. Walentiny. Numerical and analytical investigation of chemotaxis models. In *Computational Science and Its Applications – ICCSA 2018*, pages 3–18. Springer International Publishing, 2018.

[3] M. Bessemoulin-Chatard. A finite volume scheme for convection–diffusion equations with nonlinear diffusion derived from the Scharfetter–Gummel scheme. *Numerische Mathematik*, 121(4):637–670, 2012.

[4] H. M. Cheng and J. H. M. ten Thije Boonkkamp. A generalised complete flux scheme for anisotropic advection-diffusion equations. *Advances in Computational Mathematics*, 47(2):1–26, 2021.

[5] R. Eymard, J. Fuhrmann, and K. Gärtner. A finite volume scheme for nonlinear parabolic equations derived from one-dimensional local Dirichlet problems. *Numerische Mathematik*, 102(3):463–495, 2006.

[6] R. Eymard, T. Gallouët, and R. Herbin. Finite volume methods. In *Handbook of numerical analysis, Vol. VII*, Handb. Numer. Anal., VII, pages 713–1020. North-Holland, Amsterdam, 2000.

[7] P. Farrell, N. Rotundo, D. H. Doan, M. Kantner, J. Fuhrmann, and T. Koprucki. *Handbook of Optoelectronic Device Modeling and Simulation*, volume 2, chapter Chapter 50 “Drift-Diffusion Models”, pages 733–771. CRC Press Taylor & Francis Group, 2017.

[8] P. A. Farrell and E. C. Gartland Jr. On the Scharfetter-Gummel discretization for drift-diffusion continuity equations. *Computational methods for boundary and interior layers in several dimensions*, pages 51–79, 1991.

[9] H. Gajewski, H.-C. Kaiser, H. Langmach, R. Nürnberg, and R. H. Richter. Mathematical modelling and numerical simulation of semiconductor detectors. In *Mathematics — Key Technology for the Future*, pages 355–364. Springer Berlin Heidelberg, 2003.

[10] A. M. Il'in. A difference scheme for a differential equation with a small parameter multiplying the second derivative. *Matematicheskie Zametki*, 6:237–248, 1969.

[11] A. Jüngel. Numerical approximation of a drift-diffusion model for semiconductors with nonlinear diffusion. *Zeitschrift für Angewandte Mathematik und Mechanik*, 75(10):783–799, 1995.

[12] J.-G. Liu, L. Wang, and Z. Zhou. Positivity-preserving and asymptotic preserving method for 2D Keller-Segal equations. *Mathematics of Computation*, 87(311):1165–1189, 2018.

[13] D. L. Scharfetter and H. K. Gummel. Large-signal analysis of a silicon Read diode oscillator. *IEEE Transactions on Electron Devices*, 16(1):64–77, 1969.

[14] A. Schlichting and C. Seis. The Scharfetter-Gummel scheme for aggregation-diffusion equations. *IMA Journal of Numerical Analysis*, 42(3):2361–2402, 2022.

[15] J. Shen and J. Xu. Unconditionally Bound Preserving and Energy Dissipative Schemes for a Class of Keller-Segel Equations. *SIAM Journal on Numerical Analysis*, 58(3):1674–1695, 2020.

[16] J. H. M. ten Thije Boonkkamp and M. J. H. Anthonissen. The finite volume-complete flux scheme for advection-diffusion-reaction equations. *Journal of Scientific Computing*, 46(1):47–70, 2011.

[17] G. Zhou and N. Saito. Finite volume methods for a Keller-Segel system: discrete energy, error estimates and numerical blow-up analysis. *Numerische Mathematik*, 135(1):265–311, 2016.
